# Comparison of induced neurons reveals slower structural and functional maturation in humans than in apes

**DOI:** 10.1101/2020.05.21.094912

**Authors:** Maria Schörnig, Xiangchun Ju, Luise Fast, Anne Weigert, Theresa Schaffer, Sebastian Ebert, Barbara Treutlein, Nael Nadif Kasri, Benjamin Peter, Wulf Hevers, Elena Taverna

## Abstract

We generated induced excitatory sensory neurons (iNeurons, iNs) from chimpanzee, bonobo and human stem cells by expressing the transcription factor neurogenin-2 (NGN2). Single cell-RNA sequencing showed that genes involved in dendrite and synapse development are expressed earlier during iNs maturation in the chimpanzee than the human cells. In accordance, during the first two weeks of differentiation, chimpanzee and bonobo iNs showed repetitive action potentials and more spontaneous excitatory activity than human iNs, and extended neurites of higher total length. However, the axons of human iNs were slightly longer at 5 weeks of differentiation. The timing of the establishment of neuronal polarity did not differ between the species. Chimpanzee, bonobo and human neurites eventually reached the same level of structural complexity. Thus, human iNs develop slower than chimpanzee and bonobo iNs and this difference in timing likely depends on functions downstream of NGN2.

## INTRODUCTION

Differences in cognitive abilities between humans and non-human primates are thought to depend on greater numbers of neurons and more complex neural architectures in humans [1–3]. Several studies have contributed to the current understanding of the molecular and cellular uniqueness of the primate and, specifically, of the human brain. Compared to other mammals, primate brains contain a greater number of neural stem and progenitor cells that generate neurons [4–7]. In addition, some types of neurons differ structurally among primate species [8, 9]. For example, human cortical pyramidal neurons have longer and more branched dendrites than chimpanzee pyramidal neurons [10] and synaptogenesis and development of pyramidal neurons in humans and chimpanzees may be longer than in macaques [11]. Given the role of pyramidal neurons in higher cognitive functions [12], a higher degree of connectivity in human neocortex than in other primates could provide a basis for human-specific cognitive abilities [9-11,13].

Induced pluripotent stem cells (iPSCs) that can be generated from somatic cells have expanded the possibilities to compare neurogenesis among apes [14–17]. Recently, human iPSCs-derived pyramidal neurons have been found to mature slower than their chimpanzee counterparts both structurally and functionally and to end up having higher dendrite complexity and spine density [17]. In addition, upon transplantation into the mouse neocortex human neurons were shown to retain juvenile-like dendritic spines dynamics and to mature both structurally and functionally over a protracted window of time [18]. These findings are in line with previous studies that suggest that the human brain develops and matures slower than that of closely related primates [19–22].

Here, we use a direct conversion system in which chimpanzee, bonobo and human iPSCs are converted into neurons using an NGN2 overexpression system [23, 24]. Unlike what has been previously reported, we obtain a neuronal cell population containing a majority of sensory neurons. We describe their temporal differentiation in terms of gene expression, electrophysiology and dendritic arborization.

## RESULTS

### Maturation of human and ape induced neurons *in vitro*

We generated iNeurons from three chimpanzee (SandraA, JoC, ciPS01), one bonobo (BmRNA) and three human (409B2, SC102A, HmRNA) iPSC lines and from one human ESC line (H9) using forced expression of the pan-neurogenic transcription factor neurogenin 2 (NGN 2; Figure 1A) [23, 24]. We followed their differentiation using molecular, electrophysiological and morphological approaches for up to 8 weeks, as indicated in the (Figure 1A, B). Single cell RNA sequencing (scRNAseq) was performed for one chimpanzee (SandraA) and three human (409B2, SC102A1 and H9) cell lines, electrophysiology was performed using all eight cell lines and morphological analyses were done with two chimpanzee (SandraA and JoC), the bonobo (BmRNA) and three human (409B2, SC102A1 and H9) cell lines. We will refer to ape iNs for results where we combined chimpanzee and bonobo iNs for analysis.

**Figure 1.**
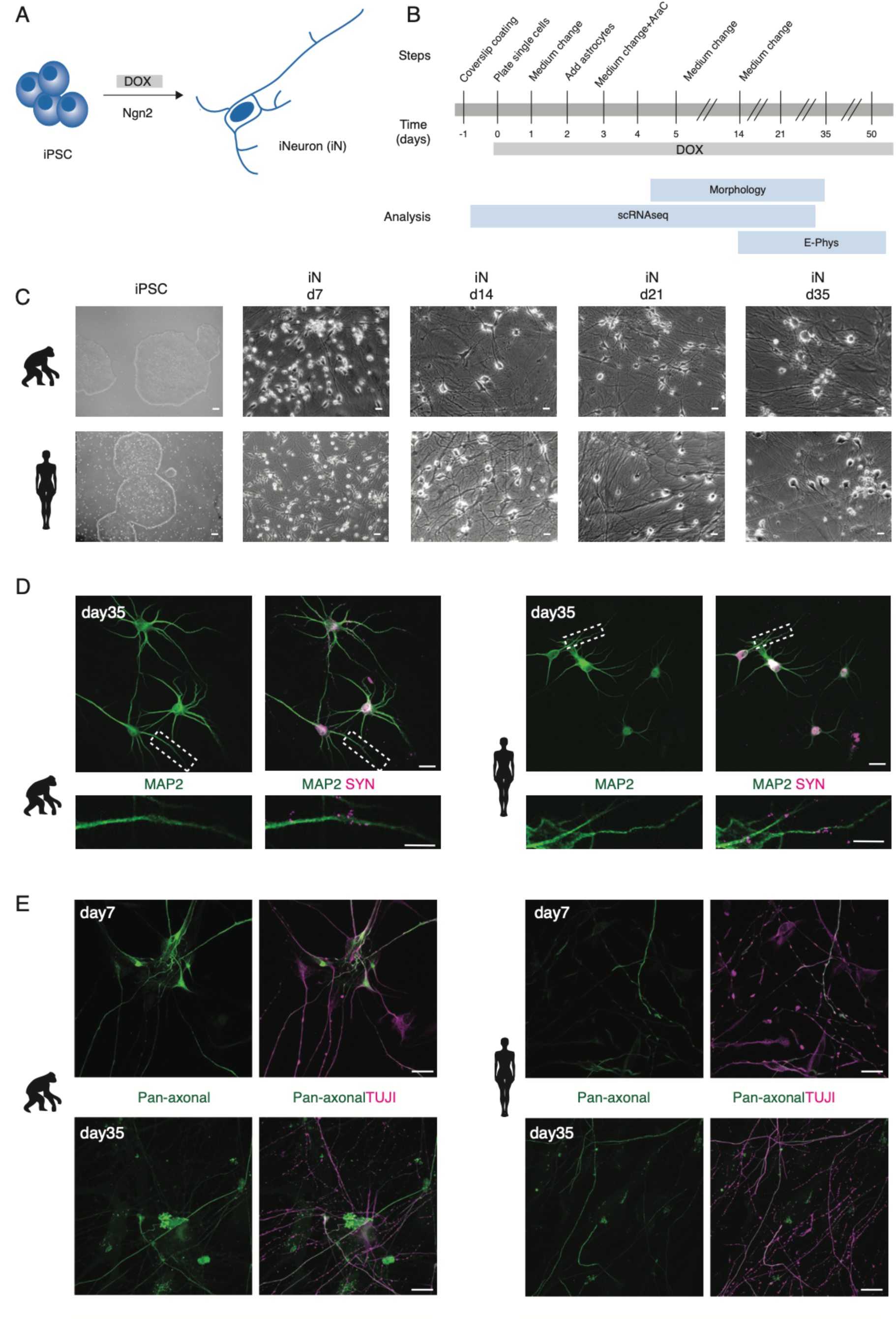
Generation of human and ape iNs. NGN2-induced iNs generated from human and ape stem cells (**A**, **B**) differentiate into mature neurons and express neuronal markers (**C**, **D, E**). **A.** iNs are generated from iPSCs/ESCs upon DOX-induced overexpression of mouse NGN2. **B.** Schematic of the experimental pipeline for iNs structural and functional analysis at different time points. The expression of NGN2 was induced at d0 by adding DOX to the culture medium. iNs were either collected for single cell RNA sequencing (scRNAseq), used for electrophysiological recordings (E-phys) or fixed for morphological analysis (morphology). Single cell RNA sequencing (scRNAseq) was performed for one chimpanzee (SandraA) and three human (409B2, SC102A1 and H9) cell lines, electrophysiology was performed using all eight cell lines and morphological analyses were done with two chimpanzee (SandraA and JoC), the bonobo (BmRNA) and three human (409B2, SC102A1 and H9) cell lines. **C.** Phase contrast images of iPSCs and (409B2) lines. Scale bars are 10 µm. **D.** Mature chimpanzee (SandraA) and human (409B2) iNs express the cytoskeletal marker MAP2 (green) and the pre-synaptic marker SYN1 (magenta). Top: low magnification view. Scale bars are 20 µm. Bottom: high magnification view, showing SYN1 puncta in juxtaposition with a MAP2-positive neurite. Scale bars are 10 µm. **E**. Distribution of TUJI (TUJI, magenta) and an axonal marker (pan-neurofilament antibody, abbreviated as Pan-Neu, green) in d7 and d35 iNs. Scale bars are 20 µm. We will refer to ape iNs for results where we combined chimpanzee and bonobo iNs for analysis.

Differentiation was characterized by a downregulation of the stem cell markers *NANOG*, *OCT4* and *SOX2* and of the neural progenitor’s markers *ID1* and *ID3* in both chimpanzee and human cells (Supplementary Figure 1), by a change in cellular morphology and by the extension of neurites (Figure 1C). Chimpanzee, bonobo and human iNs showed a neuron-like morphology at day 7 (d7) of differentiation and formed a dense network by d14. Neurites were positive for TUJI (beta-III-tubulin, a neuronal marker) starting from d3 of differentiation in apes and humans (Supplementary Figure 2).

By the end of the differentiation at d35, both ape and human cells formed networks that were positive for MAP2 (microtubule associated protein-2, marker for mature neurons) and SYN1 (synapsin-1, synaptic vesicle marker; Figure 1D). The presence of SYN1 positive punctae suggested that the iNs formed synaptic connections.

We checked for the establishment of axo-dendritic polarity by co-staining for TUJI and neurofilaments, cytoskeletal elements localized in axons (a pan-neurofilament antibody was used, abbreviated as Pan-Neu, see Supplementary Table 2 for details). At d3, TUJI largely colocalizes with neurofilaments, suggesting that the cells were not yet polarized (Supplementary Figure 3, high magnification in panel B and C). At d7, the degree of colocalization between TUJI and neurofilament markers decreased, suggesting that the iNs established axo-dendritic polarity (Figure 1E, Supplementary Figure 3). The pattern of staining of the cytoskeletal components did not differ between apes and human, suggesting that the timing of axo-dendritic polarity establishment is similar.

We next developed a sparse labeling approach that enables the tracing of individual cells in the dense connected neuronal cultures. This consisted of transfecting iNs with a GFP-encoding plasmid four days prior to fixation followed by staining with an axonal marker (Pan-Neu).The majority of iNs (25/26 cells) had a single axon, which is in line with previous findings [25]. In addition, it was always the longest neurite which was found to be positive for the axonal marker (n=26 cells, Supplementary Figure 4). Thus, in subsequent analyses, we considered the longest neurite to be the axon.

### NGN2 induces sensory neuron fates

To assess iNs identity and heterogeneity within the culture, chimpanzee (SandraA) and human (409B2) iNs were harvested at d5, 14, 28 and 35 post-induction and used for scRNAseq (10x Genomics). Two additional human cell lines (SC102A1 and H9) were harvested at d35 to assess cell heterogeneity in further detail. After data processing, across the four time points, we retained 8,786 chimpanzee iNs from the SandraA cell line and 16,921 human iNs from the human cell lines (Figure 2A-C). Unsupervised clustering analysis of the combined datasets identified 8 clusters of cells based on gene expression patterns (Figure 2B). Chimpanzee and human cells occurred in all clusters, albeit at different proportions (Figure 2A). Progenitor and neuronal cell populations were identified by *SOX2* and *MAP2* expression, respectively. The cortical markers *CUX1* and *BRN2* (Figure 2C) were expressed in both cells expressing progenitor markers and neuronal markers. Other cortical markers such as *SATB2*, *TBR1* or *CTIP2* were not detected.

**Figure 2.**
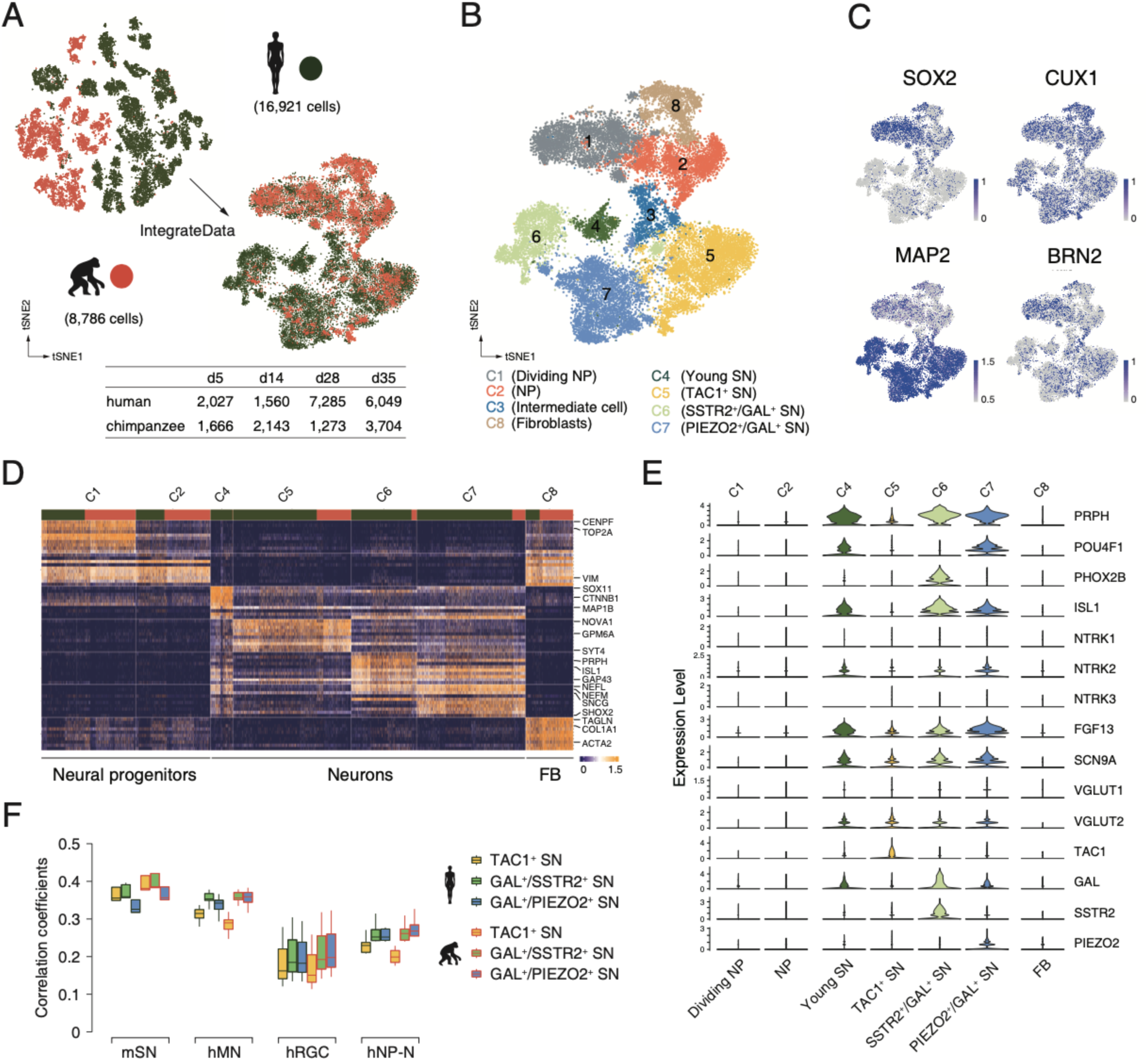
scRNAseq analysis reveals that *Nng*2-induced iNs are sensory neurons. Cell identity and heterogeneity was assessed using tSNE combined for all time points for chimpanzee (orange) and human (green) cells. **A**. Filtered cell number for chimpanzee and human cells at different time points. **B**. Identification of 8 different cell clusters by marker gene expression. NP, neural progenitor; SN, sensory neuron. **C**. Marker gene expression (blue) for progenitor cells (*SOX2*), mature neurons (*MAP2*) and cortical cells (*CUX1* and *BRN2*). Scale bars: uncorrected normalized expression. **D**. Heatmap showing scaled expression levels of top10 significantly higher expressed genes in 7 cell populations in iNs culture. The main populations we identified are fibroblasts (FB), neural progenitors and neurons. Top annotation bars: chimpanzee (orange) and human (green). **E.** The expression levels of sensory neurons markers are shown as a violin plot for neural progenitor’s clusters C1 and C2 and for neuron’s clusters C4, C5, C6 and C7. Note that the POU homeodomain transcription factors BRN3A (*POU4F1*) and Islet1 (*ISL1*) are highly expressed in a subset of young iNs. All iN clusters show high expression of growth factor FGF13 and voltage-gated sodium channel Nav1.7 (*SCN9A*) that regulate mechano-heat sensation *in vivo*. F. Pearson’s correlation of gene expressions between iNs of each species and each of mouse sensory neurons *in vivo* (mSN, GSE59739), human iPSC-derived motor neurons (hMN, GSE133764 and GSE98288), human ES-derived retinal ganglion cells (hRGC, GSE84639), and hNPC-derived cortical neurons (hNP-N, GSE142670).

The clusters include three types of neuronal cells: cells expressing progenitor markers (NP: *SOX2*^+^, *VIM*^+^, *TOP2A*^+^, *etc*.), intermediate cells (expressing both progenitor and neuronal genes) and neurons (*MAP2*, *SYT4*, *GAP43*, *etc*.). One fibroblast cluster was also identified (*COL1A1*^+^, *ACTA2*^+^ and *TAGLN*^+^, *etc*.) (Figure 2B, D).

The NGN2-induced generation of cells expressing progenitor markers and differentiated neurons is in line with previous findings [26] and with what has been reported for ASCL1-induced neuronal cell populations [27].

Based on scRNAseq, 51.5% (d5) - 21.8% (d35) chimpanzee cells and 81.8% (d5) - 66.3% (d35) human cells are identified as sensory neurons (SNs), as assessed by the expression of the sensory neurons markers *PRPH* (intermediate neurofilament peripherin), neurotrophin receptor tyrosine receptor kinase B (*TRKB*, encoded by *NTRK2*) and homeodomain transcription factors (*POU4F1*/*BRN3A*) and *PHOX2B* (Figure 2E) [28, 29], as well as by higher correlation of expressional levels to sensory neurons (GSE59739) than to other types of neurons (Figure 2F) including human iPSC-derived motor neurons (GSE133764 and GSE98288), human ES-derived retinal ganglion cells (GSE84639) and hNPC-derived cortical neurons (GSE142670). These cells express different combinations of *TAC1* (substance P), galanin (*GAL*), somatostatin receptor 2 (*SSTR2*) and piezo-type mechanosensitive ion channel component 2 (*PIEZO2*) (Figure 2B, E), normally found in somatosensory neurons sensitive to mechanical and/or thermal stimuli [30–32]. Sodium voltage-gated channel alpha subunit 9 (*SCN9A*) and fibroblast growth factor 13 (*FGF13*), which in combination selectively regulate heat nociception [33, 34], are expressed in all sensory neuron clusters (Figure 2E). Thus, NGN2 can induce the differentiation of pluripotent stem cells into nociceptive sensory neurons.

From d5 to d35, 52.1% chimpanzee and 36.9% human iNs express VGLUT2 (glutamate transporter; Figure 2E), while glutamate decarboxylases (GAD1/2, a marker for GABAergic) are not detected in either chimpanzee or human iNs, suggesting that the iNs are glutamatergic. Besides expression of sensory markers, we found that 37% (*BRN2*) to 41% (*CUX1*) of chimpanzee and 17% (*BRN2*) to 28% (*CUX1*) of human neurons express cortical markers (Supplementary Figure 5). 13% (*BRN2*) to 22% (*CUX1*) of chimpanzee and human neurons express both sensory (*PRHP*) and cortical markers (*BRN2* or *CUX1*), suggesting that a subset of neurons might be second-order sensory neurons of the cortex. In spite of the heterogeneity of the cultures, the same cell populations were found in chimpanzees and humans, albeit at different proportions, suggesting that NGN2 induces similar neuronal fates in both species.

### Transcriptional maturation of human and chimpanzee iNs

For every time point, we identified genes that are significantly higher expressed in iNs than in the cells expressing progenitor markers (NPs) at the same time point in the same species and looked for statistical enrichment of these genes among groups of genes assigned to “Biological Processes” in Gene Ontology [35]. At d5 and d14, seven to eight time more genes are higher expressed in cells classified as iNs and NPs in the chimpanzee than in the human cells and only later are numbers similar or higher in human cells (Figure 3A).

**Figure 3.**
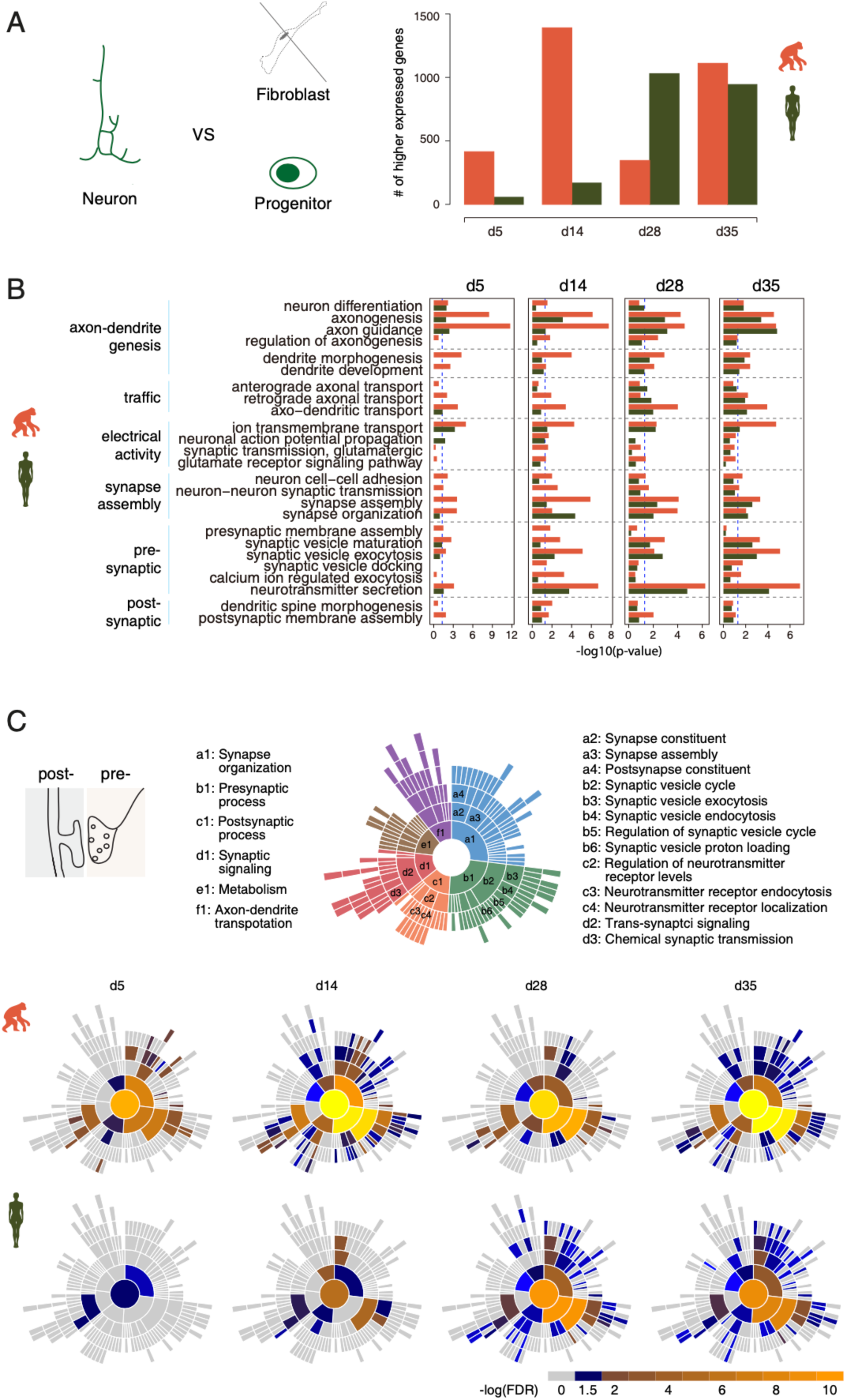
scRNAseq analysis of synaptic maturation in human and chimpanzee iNs. **A.** Higher expressed genes in human and chimpanzee iNs compared with corresponding NP of the same species at multiple time points. Left panel, schematics of cell populations for the identification of differentially expressed genes; right panel, bar plot of numbers of higher expressed genes in human and chimpanzee iNs. **B.** Categories of GO terms on biological processes (BP) enriched with higher expressed genes in chimpanzee (orange) and human (green) iNs at each time point. Reference dashed line (blue) represents p-value of 0.05 (binomial test). **C.** Sunburst plots of enriched synaptic GO terms in developing chimpanzee (top) and human (bottom) iNs. Top-left, schematic representation of synaptic parts depicted by SynGO ontology analysis. Top-right, six top-levels (a1 to f1) and selected sub-classifiers of SynGO BP terms. Bottom, the log10-transformed FDR-corrected p value per ontology term is visualized for BP. Note that the bottom BP sunburst plots are aligned with the top-right one.

Genes involved in neuron differentiation (GO: 0030182), axonogenesis (GO: 0007409) and axon guidance (GO: 0007411) are enriched (p < 0.05, binomial test) in both chimpanzee and human iNs from d5, although more strongly in the chimpanzee (Figure 3B). Genes in dendrite development (GO: 0016358) and dendrite morphogenesis (GO: 0048813) are enriched (p<0.04, binomial test) from d5 in the chimpanzee but not in humans (Figure 3B). Also, genes involved in synapse assembly (GO: 0007416) and synapse organization (GO: 0050808) are enriched from d5 in chimpanzee iNs (p < 3e-4) and become so from d14 in the human iNs (p < 0.03, binomial test) (Figure 3B). In addition, genes affecting synaptic vesicle maturation (GO: 0016188), exocytosis (GO: 0016079) and neurotransmitter secretion (GO: 0007269) are enriched from d5 in chimpanzee iNs (p < 0.02) but become enriched only from d28 in human iNs (p < 0.02, binomial test) (Figure 3B).

We also performed an enrichment analysis with neuron-expressed genes above using an expert-curated database of synaptic genes – SynGO [36]. In agreement with the results using Gene Ontology, neuron-expressed genes involved in synaptic organization, pre-synaptic process and trans-synaptic signaling (FDR < 0.05, Fisher exact test) started to be expressed in chimpanzee iNs from d5 but only from d14 and d28 in human iNs (Figure 3C). Most of the groups of genes involved in post-synaptic process and trans-synaptic signaling were not enriched in human iNs for all time points, while they were enriched in chimpanzee iNs from d14 onwards (Figure 3C and Supplementary Figure 6). Synaptic gene expression was not enriched in chimpanzee or human cells expressing progenitor markers when compared to undifferentiated iPSCs (Supplementary Figure 6B), suggesting these changes are not caused by NGN2 expression *per se* but as part of the induced neuronal maturation.

Thus, after NGN2 induction, genes associated with dendrites and synapses become prominently expressed later in human iNs than in chimpanzee iNs (Figure 3B).

### Intrinsic passive electrophysiological properties of human and ape iNs

We determined resting-membrane potential (Vrmp), cell-input resistances (Rcell), cell capacitance (Ccell) and the resulting time constant (tau = R*C; see Material and Methods for details) in 95 ape and 114 human iNs at 2 - 8 weeks after induction of NGN2 expression.

Vrmp was stable over time and similar between species (Supplementary Figure 7A; D14 - D>49, 2-way-anova (2WA) p_day_=0.23; p_spec._=0,08; p_day*spec._=0,33). Rcell decrease over time as expected from maturing neuronal cells (Supplementary Figure 7B). Although differences were not large between apes and humans, (2WA: p_day_=1,3E-8; p_spec._=0,17; p_day*spec._=0,024), chimpanzee iNs started at higher Rcell levels and decreased to about 35 % of initial values (1,2 ± 0,48 to 0,42 ± 0,18 GΩ)while human iNs started at slightly lower values and decreased to approximately 70 % at d>49 (0,79 ± 0,38 to 0,56 ± 0,27 GΩ). As expected, Ccell increased over time (Supplementary Figure 7A). While the increase occurred to a similar extent in both groups from about 20 pF to 30 pF and above, the human cells were always slightly but significantly lower than the apes. (Supplementary Figure 7A; 2WA: p_day_=1,8E-7; p_spec._=3,3E-4; p_day*spec._=0,45). The resulting time constant R*C of the cells were comparatively stable at about 15 ms and similar to values previously reported for human (or other) neuronal cells [37] (Supplementary Figure 7A). However, at early time points, the ape cells had higher values than the human cells mainly due to their high Rcel and low Ccell, respectively (2WA: p_day_=0,11; p_spec._=0,008; p_day*spec._=0,0063) (see Supplementary Table 3 for details).

Thus, the differentiation of ape and human iNs is accompanied by maturation of their basic electrical properties in a way consistent with what has been observed for other *in vitro* and *in vivo* neuronal systems [38].

### Active electrophysiological properties of human and ape iNs

Using the whole-cell current-clamp technique, we found that upon induced depolarization ape iNs showed repetitive action potentials (APs) at d10-14, with its maximal medial frequency increasing slightly over differentiation time (Figure 4A, B). In contrast, upon induced depolarization human iNs fired mainly single APs until d21, after which the ability to maintain repetitive firing became established also in the human lines (Figure 4A, B; 2WA: p_day_=0,0018; p_spec._=1,7E-7; p_day*spec._=0,56; n = 112 ape / 119 human iNs). scRNAseq analysis revealed that mRNAs for sodium, potassium and calcium channels are expressed in both chimpanzee and human iNs (Supplementary Figure 7B), with no species-specific differences.

**Figure 4.**
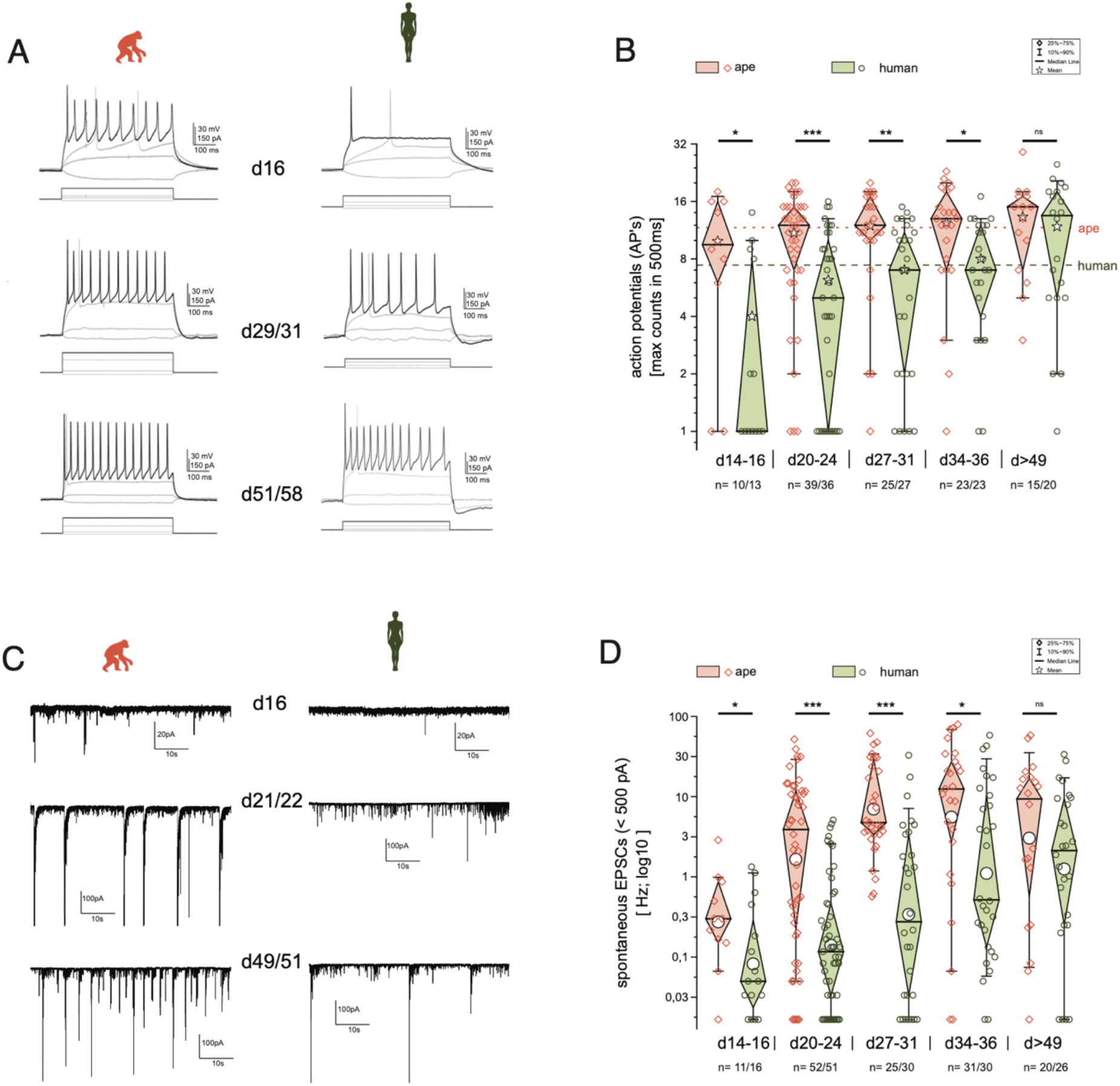
Functional development is delayed in human iNs. The development of repetitive action potentials (APs) and spontaneous excitatory post-synaptic potentials (sEPSCs) during iNs differentiation was detected using patch-clamp recordings. **A.** Voltage responses of individual cells of ape and human to depolarization by current injection at d16, d29/31 and d51/58. Given are the traces with the maximal inducible numbers of APs (black; in gray the responses to -10 to +10 pA of current injection and the trace with the first AP occurring are shown as comparisons). **B.** Boxplots of the maximal AP counts (log_2_-scale). The maximal inducible number of APs in each cell was determined and grouped according to their developmental stage representing app. 2, 3, 4, 5 and more than 7 weeks after induction of NGN2. Ape cells can maintain repetitive APs after two weeks of differentiation with a further increase during the following weeks. Human cells after two weeks of differentiation fire mainly single APs. Their ability to maintain repetitive firing developed only in the following weeks. **C.** Current responses at -80 mV over a time period of 60 s reveal spontaneous synaptic activity. Spontaneous synaptic activity is observed in ape cell lines regularly after three weeks of differentiation and cells without are rarely observed at later time points. In humans however, we observed at early differentiation only rarely spontaneous activity which came up mainly following four weeks of differentiation. **D.** Boxplots of the observed frequencies of sEPSCs in ape and human. The number of spontaneous events were counted over 60s and used for further analysis after log_10_ transformation to account for their distribution. Significance score: p<0.05 *, p<0.01 **, p<0.001 ***, Unpaired T-test with Welch Correction.

We next measured spontaneous excitatory post-synaptic currents (sEPSCs; excitatory nature verified by using kynurenic acid during recordings, data not shown) as a proxy for the establishment of functional synapses and cell-cell communication. The ape iNs developed spontaneous activity after two to three weeks of differentiation. In contrast, the majority of human iNs were silent at these times (Figure 4C-D; 2WA: p_day_=0,053; p_spec._=0,0035; p_day*spec._=0,071; n = 132 ape / 135 human iNs; see Supplementary Table 3 for details). sEPSCs became frequent in human iNs only after three to four weeks.

### Morphological heterogeneity in iN populations

Cells were fixed at d7, 14, 21 and 35 of differentiation and traced using image analysis software (Imaris, see Supplementary Table 1 for details) and quantified using custom sofware (Material and Methods). For both apes and humans, unipolar, bipolar and multipolar iNs were observed at all time points analyzed (Figure 5 A-D). For the apes and humans, 69% and 58% of neurons were multipolar, 26% and 36% bipolar, and 5% and 7% unipolar, respectively (% of all neurons from all time points). We analyzed the bipolar and multipolar iNs separately, and disregarded unipolar cells as they may be less mature, or less healthy.

**Figure 5.**
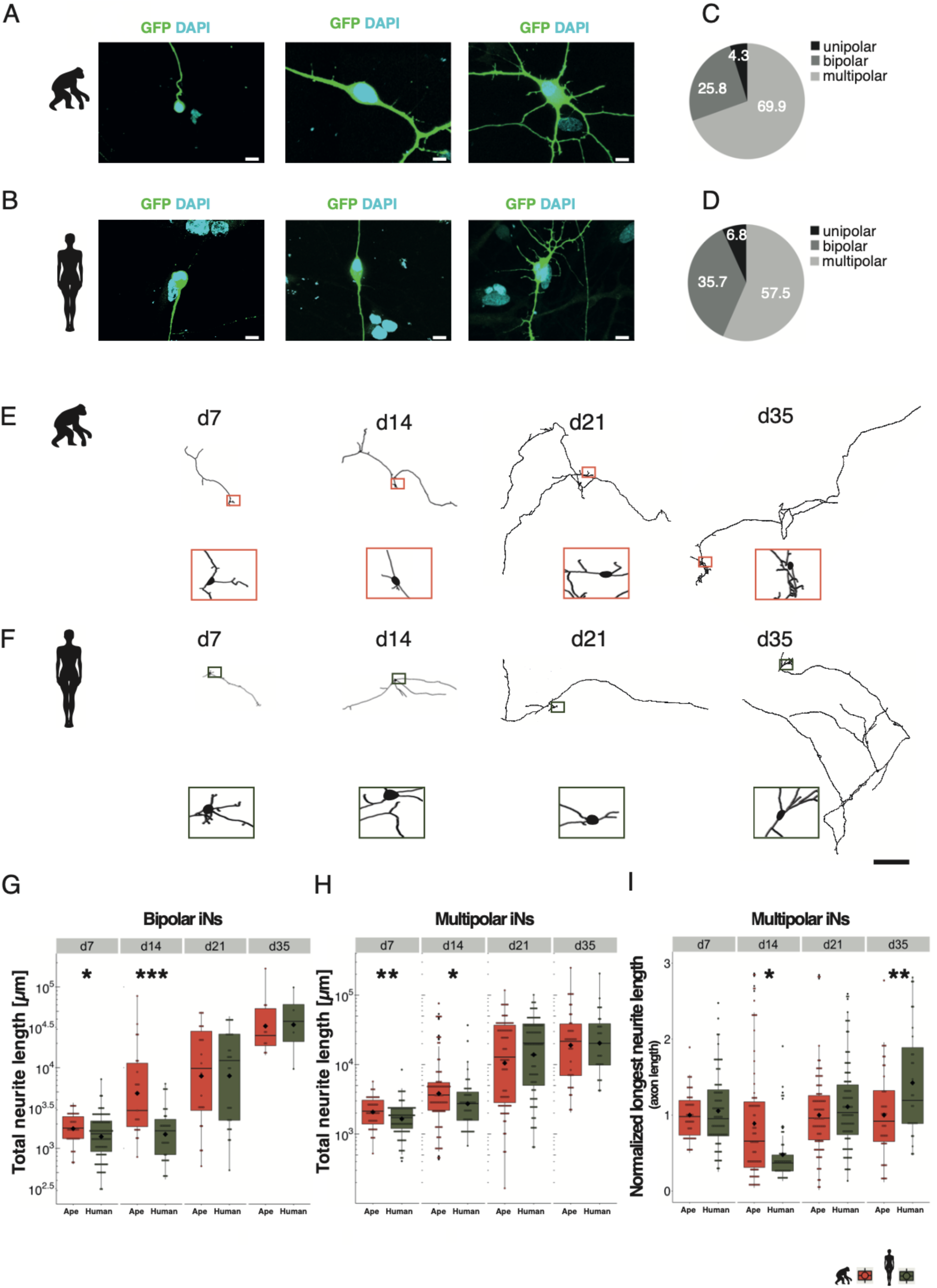
Slower morphological maturation of human iNs. iNs were lipofected four days prior fixation with a plasmid expressing cytosolic GFP and fixed at different time points. **A**, **B**. Examples of monopolar (left), bipolar (middle) and multipolar (right) iNs from chimpanzee (SandraA, **A**) and human (409B2, **B**). Cell morphology is highlighted by GFP expression (green), the nuclei are stained with DAPI (cyan). Scale bars are 20 µm. **C**, **D**. Monopolar (black), bipolar (dark grey) and multipolar (grey) iNs in ape (**C**) and human (**D**) expressed as % of total. **E, F.** Chimpanzee (**E**, SandraA, orange) and human (**F**, 409B2, green) multipolar iNs development over time. Zoom-ins show the cell body. Scale bar is 1 mm. **G, H.** Total neurite length, expressed in µm for bipolar and multipolar iNs. **G**. Apes bipolar iNs show a higher total neurite length at d7 (p=0.0278) and d14 (p=0.0003). The scale is logarithmic. **H.** Apes multipolar iNs show a higher total neurite length at d7 (p=0.0098) and d14 (p=0.0215) compared to human iNs. The scale is logarithmic. **I**. Relative length of the longest neurite (axon) in apes and human iNs. Data representation as normalized data against the mean per batch of the ape data in logarithmic scale. Ape iNs show a higher axon length at d14 (p=0.0294) compared to human iNs, according to total neurite length. Human iNs show a higher axon length at d35 (p=0.0094) compared to ape iNs. For all graphs, black rhombs represent the mean and black lines the median. Significance score: p<0.05 *, p<0.01 **, p<0.001 ***, Mann-Whitney-U-Test.

### Morphological maturation of ape and human iNs

Both bipolar and multipolar iNs from ape (multipolar iNs: Figure 5E; bipolar iNs: Supplementary Figure 8A) and human iNs (multipolar iNs: Figure 5F; bipolar iNs: Supplementary Figure 6B) show an increase in total neurite length over time. At d7 and d14 of differentiation, ape bipolar (Figure 5G, p_d7_=0.0278; p_d14_<0.001 Mann-Whitney-U-Test) and multipolar (Figure 5H p_d7_=0.0098; p_d14_=0.0215 Mann-Whitney-U-Test) iNs have neurite trees of larger total length. At d14, ape cells have more Sholl-intersections (intersections of neurites with concentric circles around the cell body) than human cells (Supplementary Figure 8E p_multipola_r=0.0469; Supplementary Figure 8F p_bipolar_<0.001 Mann-Whitney-U-Test). Thus, at early stages of maturation ape iNs have larger neurite trees than human iNs. At later time points (d21 and d35), total neurite length and number of arborizations did not differ between ape and human iNs.

Human multipolar iNs have slightly longer axons (the longest neurite) at d35 of differentiation compared to ape iNs (Figure 5I, p_d35_=0.0094 Mann-Whitney-U-Test) (see Supplementary Table 4 for detailed statistics), whereas for other neurites (excluding the longest ones), human neurons have longer total neurite length at d14 of differentiation (Supplementary Figure 8G p_multipolar_=0.0009; Supplementary Figure 8H p_bipolar_=0.447, Unpaired T-Test), but show no difference at d21 and d35.

## DISCUSSION

In this work we study the differences in neuronal maturation between apes and human by using a direct conversion protocol, in which induced neurons (iNs) are generated by forced expression of NGN2 in ape and human stem cells. We confirm and extend previous findings by showing here that NGN2 expression induces not only cortical fate [23, 24], but also sensory fate in both chimpanzee and human. Furthermore, we find that human iNs mature slower than ape iNs from a structural, transcriptional and functional point of view.

Several fundamental properties of neurons, for example the establishment of axo-dendritic polarity, onset of expression of voltage-gated channels and basic electrical properties show little or no differences between apes and humans. However, synapse-associated genes become expressed later in human neurons and electrical activity within cells (APs) and between cells (sEPSCs) appear later in the human neurons. In line with that, comparisons of human, chimpanzee and macaque *post mortem* brains have shown that expression changes of genes associated with synaptic functions and spine formation occur later in humans than in the other primates [6,20,21,39]. iNs morphological maturation follows a similar pattern where until d14 post-induction ape neurons have more complex and longer neurites, a difference that disappears by d21.

A slower neuronal development of neuronal gene expression in human than in apes has been previously described in organoids and at the organ level [16, 20]. Previous studies in cortical neurons made use of a developmental systems, in which pyramidal neurons derived from iPSC via neural progenitors (NPs) without the forced expression of NGN2 [14, 17]. The use of iNs offers us the opportunity to uncouple early developmental events such as cell cycle progression, proliferation and differentiation from the intrinsic neuronal maturation properties. Our data show that the later onset of neuron differentiation and maturation does not only occur in whole organs or complex differentiation systems but happens also in neurons as a downstream effect of NGN2-induced differentiation, strongly suggesting that the delayed maturation is a cell intrinsic property. Furthermore, our work shows that the delayed maturation occurs in sensory neurons, suggesting that delayed maturation is an intrinsic feature of human neurons, rather than a specific feature of pyramidal neurons. Of note, sensory neurons are interesting from an evolutionary point of view, as the development and evolution of working memory in human is linked to a higher integration of sensory functions in the human prefrontal cortex [3,40–42]

There are several possible consequences the delayed development and maturation of human neurons may have for the organization and function in the adult brain. The later onset of neurite outgrowth shown here and by others [14,17,39,43] has been speculated to result in more complex dendritic trees [10,13,17]. Interestingly, at d35, we find that the axons of the human multipolar neurons are slightly longer than in the ape neurons. Further work is required to understand if longer axons allow neurons to develop further and establish more contacts with other neurons. Of note, longer axons would obviously make contacts over greater distances in the brain possible. Since the human brain is three times bigger than the ape brains, in the future it will be interesting to investigate if axons may be longer in human than in ape brains, and if that may affect the extent of neuronal projections in human brains.

## MATERIAL AND METHODS

### Generation of rtTA/*Ngn*2-positive stem cell lines

For this study we used three human (hiPS409-B2, SC102A1, HmRNA), three chimpanzee (SandraA, JoC, ciPS01) and one bonobo (BmRNA) iPS cell lines and one additional human ES cell line (H9). The human iPSC lines hiPS409-B2 and SC102A1 were purchased from the Riken BRC Cellbank and System Biosciences, respectively. The human iPSCs line HmRNA (generated in this study) was reprogrammed from human dermal fibroblasts using the StemMACS mRNA transfection kit. The cell line was validated for pluripotency markers by immunohistochemical staining using the Human Pluripotent Stem Cell 3-Colour Immunohistochemistry Kit and were differentiated into the three different germ layers using the Human Pluripotent Stem Cell Functional Identification kit and StemMACS Trilineage Differentiation Kit. Karyotyping was carried out using Giemsa banding at the Stem Cell Engineering facility, a core facility of CMCB at Technische Universität Dresden. Karyotypes were found to be normal. The human ES cell line H9 was purchased from WiCell. The chimpanzee iPSC lines SandraA and JoC as well as the bonobo iPSCs line BmRNA were generated in a previous study [16]. The chimpanzee iPSCs ciPS01 line was provided by the Max-Delbrück-Centrum für Molekulare Medizin, Berlin.

The rtTA/*Ngn*2-positive iPSCs/ESCs hiPS409-B2_*Ngn*2, SandraA_*Ngn*2, BmRNA_*Ngn*2, H9_*Ngn*2, SC102A1_*Ngn*2, HmRNA_*Ngn2,* ciPS01*_Ngn2* and JoC_*Ngn*2 were generated using lentiviral vectors to stably integrate the transgenes into the genome of the stem cells as previously described by Frega *et al.* [24].

We used the human lines hiPS409-B2_*Ngn*2, H9_*Ngn*2 and SC102A1_*Ngn*2, the chimpanzee lines JoC_*Ngn*2 and SandraA_*Ngn*2 and the bonobo line BmRNA_*Ngn*2 for the electrophysiological and morphological experiments. For the electrophysiological experiments we used additionally the human iPS cell line HmRNA_*Ngn2* and the chimpanzee cell line ciPS01_*Ngn2*. For the scRNAseq experiments we used the hiPS409-B2_*Ngn*2 and SandraA_*Ngn*2 lines and in addition for d35 the human lines H9_*Ngn*2 and SC102A1_*Ngn*2.

### Culturing of stem cell lines

Human, chimpanzee and bonobo rtTA/*Ngn*2-positive stem cells were cultured in 6-well cell culture plates coated with Matrigel (1:100 dilution in Knockout DMEM/F12) and maintained in mTeSR™1 with supplement and antibiotics (dilution Pen/Strep 1:200, G418 1:1000 and Puromycin 1:2000) at 37 °C and 5 % CO_2_. The medium was refreshed daily.

To maintain the culture, stem cell colonies with a round and densely packed morphology were either picked and seeded into wells containing fresh medium or passaged once they reached a confluency of approximately 80 %. For passaging, the cells were washed with DPBS before incubation with 1 ml EDTA (1:1000 dilution in PBS) for 5 min at 37 °C to detach the cells from the culture dish. EDTA was aspirated and cells were detached with 1 ml medium. After splitting, the cell suspension was transferred into new wells containing fresh mTeSR™1 with supplement. Rock Inhibitor (1:1000 dilution) was added for the first 24 h of the culture. Our cultures were regularly controlled for mycoplasma.

### Cryopreservation of stem cells

Cells were collected and prepared for freezing by washing them with DPBS twice before incubation with 1 ml TrypLE™ Express for 5 min at 37 °C. Cells were washed from the culture dish with 3 ml DMEM/F12 and centrifuged at 200 x g for 5 min. The pellet was resuspended in 500 µl mFreSR™, transferred into a cryogenic vial and gradually frozen to – 80 °C in an isopropanol filled freezing box.

To recover the cells, the vial was quickly thawed in a water bath at 37 °C. The cell suspension was transferred into 5 ml DMEM/F12 and centrifuged at 200 x g for 5 min. The cell pellet was resuspended in 2 ml medium and transferred into fresh wells containing mTeSR™1 with supplement and Rock Inhibitor (1:1000 dilution) was added for the first 24 h of culture.

### Differentiation of rTTA/*Ngn*2-positive stem cells to iNeurons

Human, chimpanzee and bonobo stem cells were differentiated into induced Neurons (iNs) according to Frega *et al*., 2017 [24]. Briefly, we plated iPSCs or ESCs as single cells and initiated the neuronal differentiation by doxycycline-induction of NGN2. After two days of culturing we added primary cortical rat astrocytes to the system. One day after astrocyte plating we stopped proliferation of undifferentiated stem cells and proliferating astrocytes by supplementing the growth medium with 2 µM cytosine arabinoside (Ara-C). iNs were kept for two to eight weeks in culture with 50% medium changed every two to three days (Figure 1B).

### Single cell transcriptomic analysis

#### Single cell RNA-seq data generation

For each cell line, an entire 6 well plate was plated with 120.000 to 150.000 iPSCs and 100.000 astrocytes/well. The cells were differentiated into iNs according to the abovementioned protocol. On d5, d14, d28 and d35 the cells were dissociated into single cells to perform a 10x analysis. All reagents were filtered through a 0.22 µm filter. Each well was washed with DPBS before the cells were incubated for 5 min in 1 ml EDTA 1 % accutase/well. Using 1 ml pipette tip, cells were carefully detached from the plate by continuous rinsing to obtain a cell suspension enriched for iNs. All wells of one cell line were pooled and centrifuged for 5 min at 300 g. Then, the supernatant was discarded and the pellet was resuspended in 1 ml accutase with 0.3 µl DNAse (2000 U/µl) for 5 min. The cell pellet was washed 2-3 times in 5 ml Neurobasal medium supplemented with 10% FCS followed by centrifugation for 5 min at 300 g. Then, the pellet was pipetted 20 times until complete resuspension in 20-00 µl Neurobasal medium supplemented with 10% FCS and filtered through a 30 µm filter. Finally, we centrifuged cells on a Percoll gradient for 5 min at 300g to enrich for healthy cells and remove debris. The Percoll gradient consisted of three layers. Lowe layer: 4 ml wash medium and 1 ml Percoll solution; middle layer: 4 ml wash medium and 0.75 ml Percoll solution; upper layer: 4 ml wash medium and 0.5 ml Percoll solution. Cells were analyzed using Trypan Blue assay and counted with the automated cell counter Countess (Thermo Fisher) and diluted to yield a concentration of 500 to 1000 cells/ml to obtain approximately 3000 cells per lane of a 10X microfluidic chip device. Human and chimpanzee cells were pooled for the 10X experiments and separated computationally afterwards.

Single cell gene expression libraries were generated using the 10X Chromium Single Cell 3’ v2 Kit following the manufacturer’s instructions. Quantification and quality control of libraries was performed using a High Sensitivity DNA chip for Agilent’s Bioanalyzer and sequenced on a HiSeq2500 in Rapid sequencing mode.

#### Data processing

Chromium single-cell RNA-seq (scRNAseq) output was processed using Cell Ranger pipelines. Raw base call (BCL) files were first demultiplexed into FASTQ files with Cellranger mkfastq. Raw reads were aligned to human hg19 and mouse mm10 reference genomes to separate primate cells versus rat astrocytes. After removal of rat astrocyte cells and primate-rat multiplets, scRNAseq runs that included both a chimpanzee and a human batch were computationally separated. FASTQ files of a published scRNA-seq dataset from cerebral organoids containing the same lines as used in this study were processed equally with Cellranger (v2.1.0) count using GRCh38 as a reference. The resulting BAM files were processed to the VCF format using SAMtools 1.3.1 and BAMtools 1.4. The obtained polymorphisms were used to demultiplex the cells using demuxlet [44]. We retained genes and single cells for downstream analysis also based on following quality measures: mean count in at least one batch of cells larger than 0.1; We removed cells with less than 500 or more than 6000 genes detected and cells in which more than 5% of the transcripts accounted for mitochondrial genes were. After filtering, 25,707 cells on culturing day 5, day14, day 28 and day 35 were retained, and 1,993 genes and 7,051 UMIs were detected in each single cell on average. For human and chimpanzee iPSCs, 3,329 cells were retained after filtering, and 2,140 genes and 7,651 raw reads were detected in each single cell on average.

#### Identification of neuronal cells and differentially expressed genes

Data normalization, scaling and variable feature finding were performed using “SCTranform” function of *Seurat* R package (v3.0.0) [45]. Multiple batches of single-cell datasets were integrated with “FindIntegrationAnchors” and “IntegrateData” functions of *Seurat* R package with the first 20 components were used for finding the anchors [45], and 1,158 highly variable genes were obtained, which were used to perform principle component analysis (PCA). The first 27 PCs were estimated significantly enriched with variable genes based on the percentage of variance explained by each one with the “ElbowPlot” function in the *Seurat* R package and were selected as the significant components for *t*-SNE analysis. We performed clustering of single cells with the parameter *resolution* to 0.6 using “FindClusters” function in *Seurat*. Clusters with similar marker gene expressions were merged manually into one. Neuronal cells and progenitors were identified according to their expressions of corresponding marker genes: progenitor markers (*SOX2, VIM, ID3, MKI67, TOP2A, CENPF*) [46], pan-neuron markers (*MAP2, SYT4, SYT1, STMN2, GAP43, DCX, NEFM, VGLUT2*)) [47, 48], sensory neuron genes (*PRPH*, *ISL1*, *BRN3A*, *PHOX2B*, *TAC1*, *GAL*, *SSTR2*, *PIEZO2*, *FGF3*, *SCN9A*, *NTRK2*) [28, 29] and fibroblast cluster markers (*COL1A1, COL1A2, MYL9, ACTA2, TAGLN*) [49].

Differentially expressed genes (DEGs) between neuronal cells and neural progenitors of each species at each time point were identified by using the “FindMarkers” function in the *Seurat* R package with the cut-offs of adjusted p value less than 0.05 and log2 transformed fold change larger than 0.25. We here focused only on DEGs having higher levels in neuronal cells (neuron-expressed genes).

To compare gene expressions before and after NGN2 induction in differentiated (neurons and progenitors) and iPSCs, expressional datasets were integrated with the same procedure as described above. New cell identities of chimpanzee and human iPSCs, neurons and progenitors were set, using the “Idents” function in the *Seurat* R package, and then DEGs between cell identities were identified with the “FindMarkers” function in *Seurat*.

#### Gene ontology enrichment analysis

DEGs (neuron-expressed genes) of each species were used to identify over-represented biological processes (BPs) of gene ontologies (GOs) and the genes detected by scRNAseq in chimpanzee or human neurons and progenitors were used as the background, respectively. The human GO database on biological process was download with the “loadGOTerms” function in the *goSTAG* R package including 18,446 genes in 3,599 GO categories of biological processes [35]. The exact binomial test was used to identify categories of biological processes with an enrichment of DEGs (neuron-expressed genes) with a high fraction compared to the fraction of detected category genes in the background list. The binomial test was performed using the “binom.test” function in the R package. The significantly enriched GO categories were selected based on the p values being less than 0.05. To visualize how DEGs (neuron-expressed genes) regulate synaptic functions at each time point, we perform GO enrichment analysis with the online *SynGO* annotations including 179 synaptic processes based on published, expert-curated evidence [36]. Significantly enriched synaptic processes were retained with the FDR less than 0.05.

### Electrophysiology

iNeurons were grown onto poly-L-ornithin and mouse laminin-coated coverslips (12 mm diameter, Kleinfeld, Germany). Individual coverslips were carefully transferred to a recording chamber under an upright microscope (Olympus BXW-51, Hamburg, Germany) and were perfused (2-3 ml/min) with an artificial cerebrospinal fluid (aCSF; in mM: 100 NaCl, 3.5 KCl, 1 MgCl_2_, 2 CaCl_2_, 30 NaHCO_3_, 1.25 NaH_2_PO_4_, 10 glucose) continuously bubbled with carbogen (95% O_2_ and 5% CO_2_) to maintain a pH < 7.4. Recordings of single cells were done in the whole-cell configuration of the patch-clamp technique with glass electrodes (4.5 - 5.5 MΩ; oD/iD, borosilicate, Hilgenberg, Germany), pulled to tip resistances of ∼ 5.5 MO (Sutter, P97) and filled with an internal solution containing 130 mM K-gluconate, 10 mM NaCl, 4 mM Mg-ATP, 0.5 mM GTP, 10 mM HEPES and 0.05 mM EGTA (adjusted to pH = 7.3; filtered at 0,2 um). The liquid junction potential (15 mV) was corrected online. Intra- and extracellular solutions were adjusted to 270 and 280 mOsm respectively. This osmolarity closely matched the cell-culture medium and allowed for long-time recordings with stable series-resistances (Rs, typically < 15 MΩ). Recordings were carried out at room temperature (22-24°C) using an EPC-10 amplifier (HEKA, Lambrecht, Germany). Acquisition and analysis were performed using Patch- and Fitmaster software (Ver. 2.9x, HEKA) in conjunction with Origin (Ver 2018-2019b, OriginLab, Northampton, MA, USA). Recordings were digitized at 10 or 20 kHz and done in either the voltage- or current-clamp configuration. In the voltage clamp configuration 50% Rs-compensation was applied routinely, in the current-clamp the bridge-compensation was used to compensate any voltage-drop across the series resistances.

#### Recordings

The measured the following basic properties: (i) Vrmp or membrane potential, determined by the concentration of ions on both sides of the membrane, the membrane’s permeability and activity of ion-pumps (ii) Rcell or cell-input resistances, influenced by the cell size and the number of membrane proteins e.g. ion-channels & transporters (iii), Ccell or cell capacitance, reflects the membranes characteristic to act as a capacitor; as the capacitance influences the impedance -the resistance to AC currents-it influences the propagation of fast changing signals e.g. APs (iv) tau = R*C or resulting time constant, a measure of the cell response to stimuli.

The basic properties of the iNs as cell-input resistances (Rcell) and whole-cell capacitances (Ccell) were obtained from the software and verified using the following two protocols. First, we recorded four successive uncompensated voltage steps from -80 to - 95/-85mV/-75/-65mV (20kHz sampling rate) to obtain Ccell from the initial fast capacitive decay and Rcell from the plateau current (details see analysis). Second, we used a ramp-protocol (from -90 mV (200 ms) rising to +60 mV within 800 ms; back to -90 mV) to verify the presence of voltage activated currents and to determine Rcell from the current slope and the resting membrane potential (Vrmp) from the zero-current crossing of the voltage ramp. Thereafter we recorded EPSCs at -80 mV for 60 s without pharmacological interference (10 khz sampling rate, filtered at 2.5 kHz). In few recordings we applied 50 µM kynurenic acid to verify their glutamatergic identity. We then switched to the current-clamp configuration and measured the induction of action potentials by depolarization. We determined the current ΔI that induced app. 5mV change at rest (app. -80mV). We then hold the cell at app. 90 mV to remove any inactivation of voltage gated ion-channels and recorded 20 current steps (500 ms long, 2 hyper-, 18 depolarizing) with changing current amplitudes as determined previously (ΔI).

#### Analysis

To obtain Ccell from the first protocol the initial 500 us of the fast-capacitive decay were fitted with an exponential function. The obtained time-constant tau allowed determination of Ccell (C = Rs / tau). In addition, Ccell was determined using the RC compensation routine from the Patchmaster software, once automatically and once manually under visual control. The plateau currents after the transient decay (10 ms mean after 20 ms) allowed determination of Rcell (Rcell = ΔU / ΔI). A second value we obtained from the ramp-protocol using the current slope between 75 to 65 mV (Rcell = ΔU / ΔI) and a third value from the amplifier of the Patchmaster software. Normally all three values for Ccell and for Rcell were in good agreement (<10%) and were averaged. In case of disagreements the traces were reanalyzed and the accurate value determined by hand. The resting membrane potential (Vrmp) was obtained from the zero-current crossing of the voltage ramp. In some cases where a low subthreshold activation of voltage-gated sodium currents hyperpolarized the membrane close to and before the zero-current crossing, the resting potential was estimated using a linear fit to the slope between -75 and -65 mV and determining its zero crossing. Action potentials in the current clamp protocols were determined as positive crossings at -20mV. The maximal numbers obtained during these steps were used for analysis. EPSCs traces were exported for further analysis to the program NeuroExpress (NeXT, v. 19.n; Attila Szücs, UCSD San Diego) via an Igor binary format. There we used the *“Triangular template”* with the following settings for an analysis of single excitatory events: estimated noise: 14 pA, max EPSC rise-time: 1.5 ms, max EPSC fall time: 6 ms, max EPSC amplitude: 500 pA.

### Lipofection of iNeurons

To generate GFP expressing iNs, cells were lipofected four days before fixation using the Lipofectamine 3000 Transfection Kit from Invitrogen. Briefly, 5 µl OptiMEM were mixed with 0.75 µl Lipofectamine and then added to 25 µl OptiMEM containing 0.1 µg GFP pMAX plasmid (Lonza) and 1 µl P3000 reagent. After incubation for 15 min, 50 µl lipofection mix were added per coverslip and the iNs were kept in culture until fixation.

### Immunostaining of iNeurons

#### Preparation of paraformaldehyde fixative

To prepare 100 ml 4% paraformaldehyde, all steps were performed under the hood. 4 g PFA were added to 40 ml distilled water and were heated to exactly 60 °C while stirring. Then, 1 N NaOH was added dropwise until the solution cleared. 4 g sucrose and 50 ml 240 mM Na-Phosphate buffer (pH 7.4) were added to the mixture. After cooling, the pH of the solution was checked and if necessary, adapted to pH 7.4 by adding HCl. The solution was brought to a final volume of 100 ml by adding bidistilled H_2_O and then filtered through a 0.22 µm filter. 5-10 ml aliquots were stored at -20 °C.

#### Fixation of GFP-labelled iNeurons

GFP-pmax-lipofected iNs were fixed at day 7, 14, 21 and 35 of differentiation. The coverslips were transferred with tweezers from the 24-well cell culture plate into a tissue culture dish (35 mm) containing 1 ml PBS. 1 ml 4% PFA (pre-warmed to 37 °C) was added to the dish under a hood and the cells were incubated for 8 min. The coverslips were washed with PBS three times and stored in PBS at 4 °C. The coverslips were blind after the fixation step, before the start of the immunostaining.

#### Quenching and immunostaining of GFP-labelled iNeurons

*Analysis of GFP-labelled iNeurons direct fluorescence*. PBS was aspirated and the coverslips were incubated in 1 ml 0.2 M Glycine buffer for 30 min. After three quick washed with immunofluorescence buffer (IF-buffer: 120 mM phosphate buffer cointaining 0.2% gelatin and 0.05% Triton X-100) the coverslips were mounted on slides with Mowiol 4-88 and DAPI (1:1000 dilution) and stored at 4 °C.

*Immunostaining of GFP-labelled iNeurons*. iNs were permeabilized with 0.05% Triton X-100 in PBS for 10 min and quenched in 1 ml 0.2 N Glycine buffer for 30 min at room temperature. The coverslips were washed with PBS and incubated with 100 µl of the respective primary antibody diluted in IF-buffer (see Supplementary table for the list of Abs). The coverslips were washed with IF-buffer five times for 5 min and then incubated with 100 µl of the respective secondary antibody diluted in IF-buffer containing DAPI (dilution 1:1000). After five washes with IF-buffer for 5 min and three quick washed with PBS, the coverslips were mounted on slides with Mowiol 4-88 and stored at 4 °C.

### Image Acquisition

iNs were acquired as confocal Z-stacks. iNs were imaged using an Olympus FV1200 confocal microscope equipped with a 40 x oil immersion objective (optical section thickness: 1.028 µm, distance between consecutive optical sections: 0.4 µm) or a Zeiss LSM 780 NLO 2 photon upright microscope equipped with a 63x Neofluar immersion objective (optical section thickness: 0.8 µm, distance between consecutive optical sections: 0.6 µm). The acquisition of day 35 data set was carried on using a spinning disk Andor IX 83, inverted stand, equipped with a 60x oil immersion objective. We acquired three-dimensional (Z-stack) tile scans with a number of z-sections ranging from 5 to 30 depending on the cell. Single tiles were 1024 x 1024 pixels. The following lasers were used: 405 nm for DAPI, 488 nm for GFP and 561 nm for RFP/Alexa-555 were used. The acquisition experiments were performed in blind.

### Image quantification

*Quantification of neuronal morphology*. We quantified GFP-expressing iNs at d7, 14, 21 and 35 of differentiation (see Supplementary Table 1 for a summary of the number of cells traced). Three-dimensional confocal z-stacks were traced and analyzed using Imaris9.5 [50]. We used a manual tracing approach with a neurite’s diameter for of 0.4 µm. For quantification, we exported Sholl intersections and .hoc files, as they contain information about both the neurite length and the branching pattern. HOC files were analyzed using a custom script [51]. Statistical analyses were performed using R version 3.4.4. Experiments were unblinded only after running the complete quantifications. Note that the increase in neurite length is batch-dependent for both bipolar and multipolar cells (a batch is defined as one experiment of differentiation of stem cells into iNs; bipolar cells: F(1,220)= 13.1790, p_batch_<0.0012WayAnova; multipolar F(1,454)= 17.2302, p_batch_<0.001). In Figure 5 and Supplementary Figure 8 we normalized for the batch effect by dividing the data for length of the longest neurite by the mean of the ape data per batch. *Quantification of TuJI signal.* In Supplementary Figure 2, the images of the time course of TUJI expression were acquired sequentially using the same settings during one session at the Zeiss LSM 780 NLO microscope. For each image, we use FiJI and quantify the signal in an area containing only neurites (but not cell bodies). The signal is expressed as arbitrary units (AU).

## ACKNOWLEDGEMENTS

The authors thank Svante Pääbo for the continuous support and discussion on the project. We thank Kathrin Köhler, Linda Dombrowski, Katrin Linda, Teun Klein Gunnewiek, Noor Smal and Malgorzata Santel for excellent technical help, support and advice, the Tchimpounga Sanctuary for support with usage of the JoC iPSC line and the Max-Delbrück-Centrum für Molekulare Medizin, Berlin for providing the chimpanzee iPSC line Chimp male iPSC Sendai CL5 (ciPS01), the light microscopy facility at the MPI-CBG in Dresden, especially Sebastian Bundschuh and Britta Schroth-Diez, for help and assistance with confocal microscopy. Experiments and generation of NGN2-inducible stem cell lines were in part performed at Radboudumc, Nijmegen, The Netherlands. This work was supported by the Max Planck Society.

We thank Svante Pääbo, Takashi Namba, Nereo Kalebic and Sabina Kanton for comments and discussion on the manuscript.

## AUTHOR CONTRIBUTIONS

MS generated *Ngn*2/rtTA positive stem cell lines and grew iNs cultures with assistance from LF, AW and TS. AW generated and characterized the human iPS cell lines HmRNA and HmRNA_*Ngn2* and the chimpanzee iPS cell line ciPS01_*Ngn2*. MS, ET and LF performed immunohistochemical staining and microscopy. BP and MS wrote the custom script for morphological analyses. MS, LF and AW performed morphological analyses. MS, SE, LF and AW performed single cell RNAseq experiments. XJ and SE analyzed single cell RNAseq data. WH performed and analyzed electrophysiological experiments. NNK provided information and material relevant for the iNs generation and interpretation of the data. MS, ET and BT designed the study. MS, XJ, WH and ET wrote the manuscript with support from all authors.

