## Supplementary Figures for "Comparison of induced neurons reveals slower structural and functional maturation in humans than in apes"

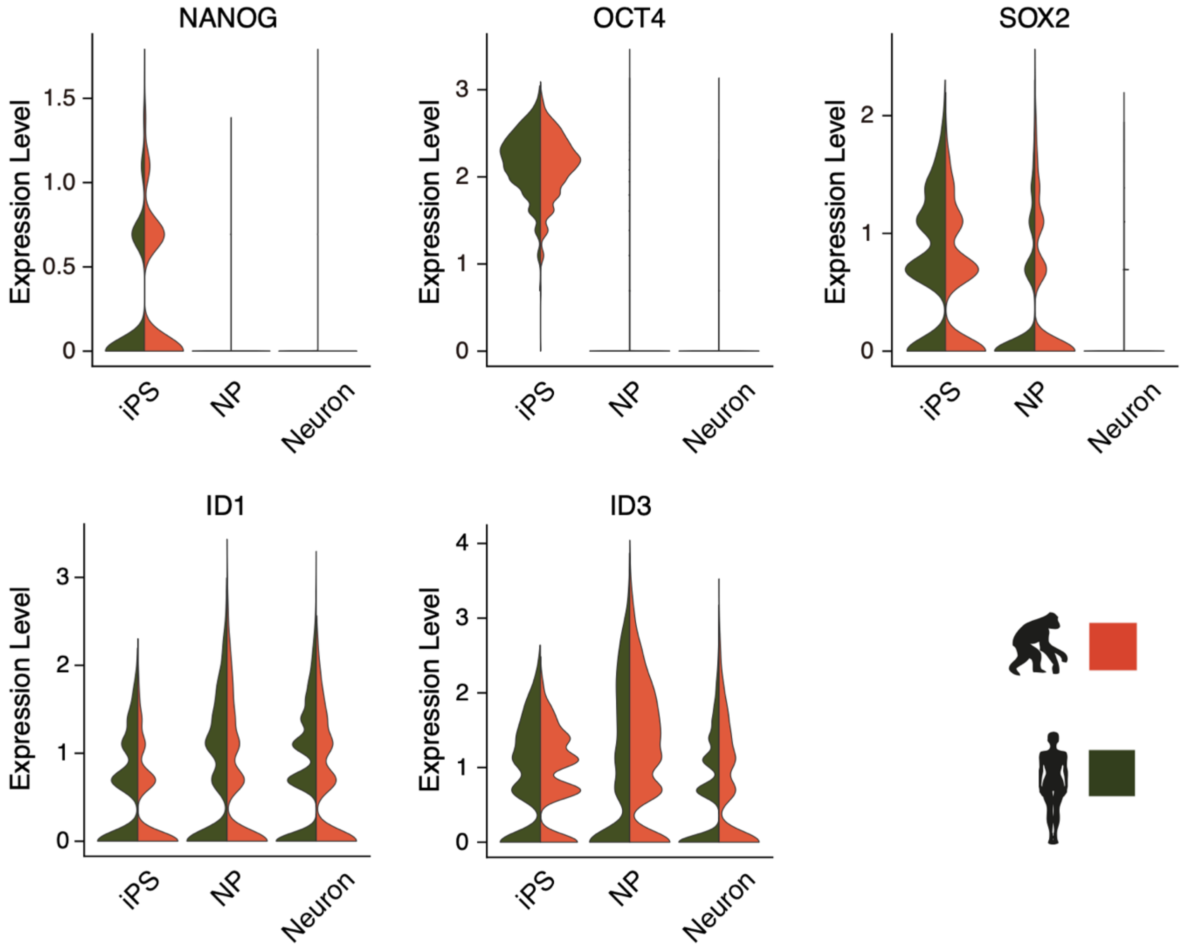


**Supplementary Figure 1. Downregulation of stem cell markers in iNs.** The violin plots show the expression in iPSCs, neural progenitors (NP) and neurons of three stem cell marker genes (*NANOG*, *OCT4* and *SOX2*) and two neural progenitors' markers (*ID1, ID3*) in chimpanzee and human iNs. All stem cell genes show a reduction in their expression at the transition from iPSCs to NPs.

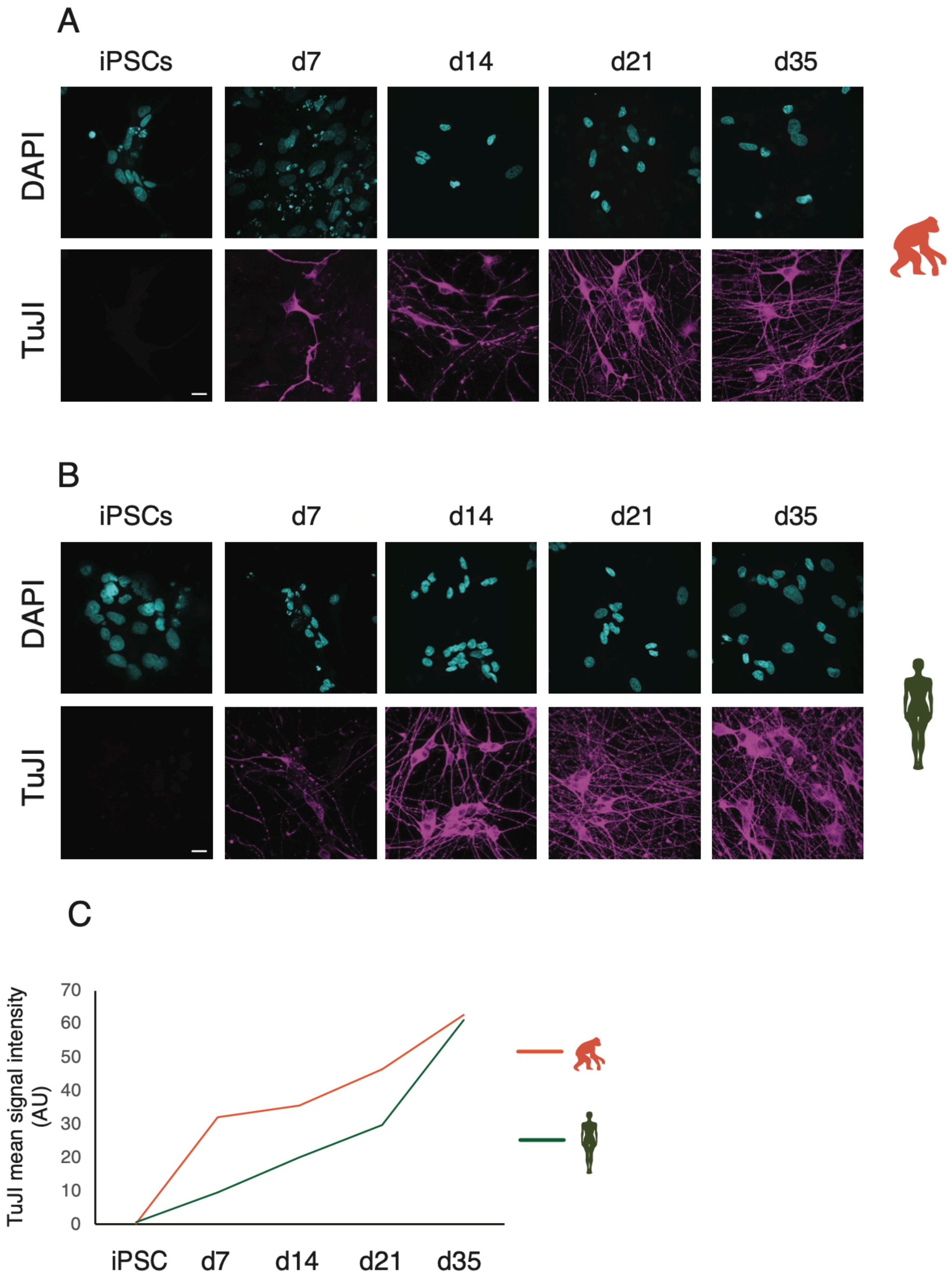
A
iPSCs
d7
d14
d21
d35
DAPI
TuJI
B
iPSCs
d7
d14
d21
d35
DAPI
TuJI
C
TuJI mean signal intensity (AU)
70
60
50
40
30
20
10
0
iPSC
d7
d14
d21
d35

**Supplementary Figure 2. iNs express neuronal markers.** Time course of TUJI expression during differentiation of iNs from iPSCs in chimpanzee (**A**, SandraA) and human (**B**, 409B2). Magenta: TUJI; cyan: DAPI. Scale bars are 20 $\mu$m. **C**. Quantification of TUJI signal intensity in chimpanzee (orange) and human (green) iNs. The intensity of the signal was quantified using FiJI and is expressed as AU (arbitrary unit; see Material and Methods).

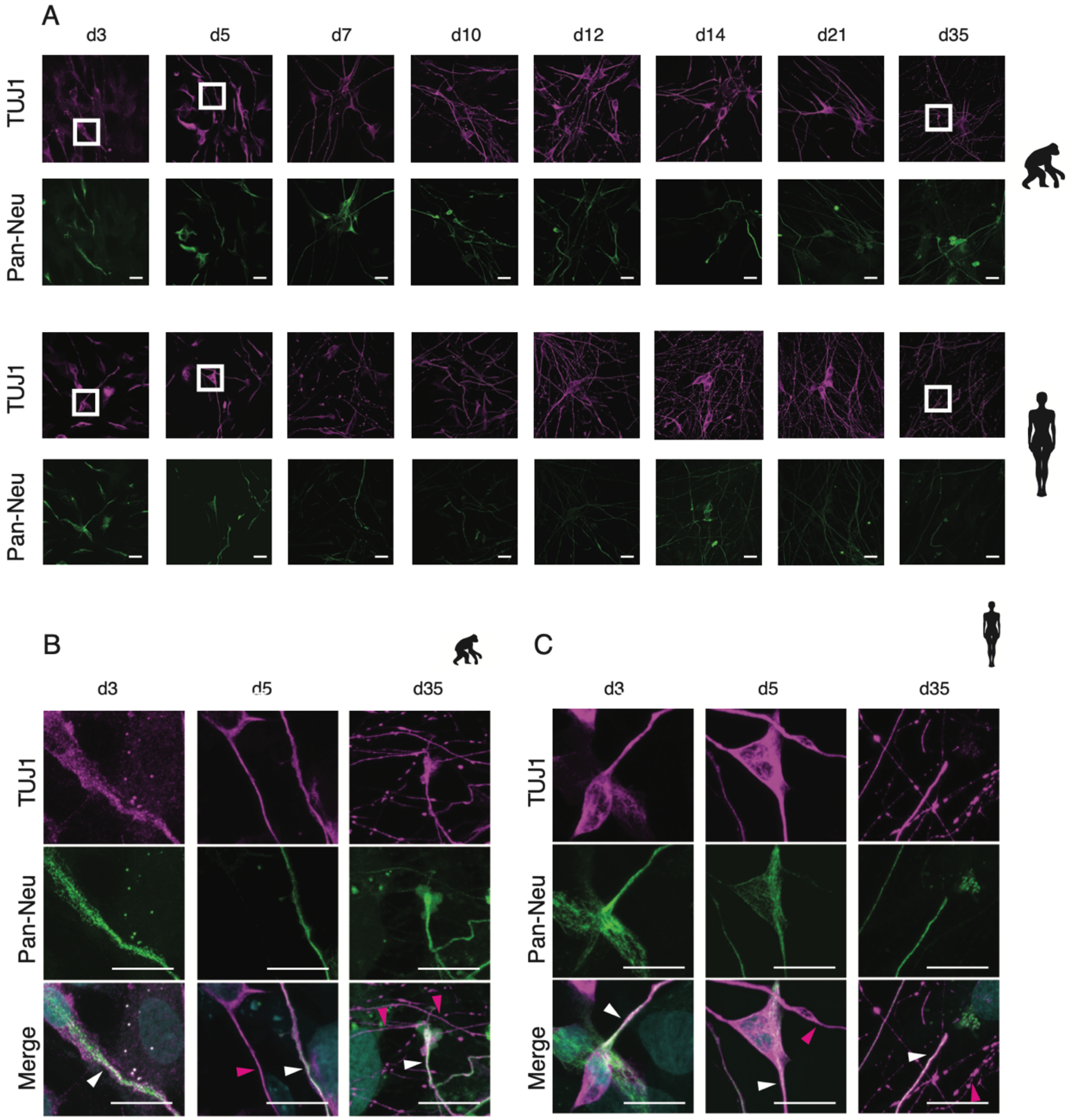


**Supplementary Figure 3. Polarity establishment in ape and human iNs.** Axo-dendritic polarity is established around d7 in chimpanzee (SandraA) and human (409B2) iNs. **A**. Distribution of TUJI (TUJI, magenta) and an axonal marker (pan-neurofilament antibody, abbreviated as Pan-Neu, green) during iNs differentiation. Scale bars are 20 $\mu$m. **B**. High magnification of the areas highlighted with a white box in **A.**

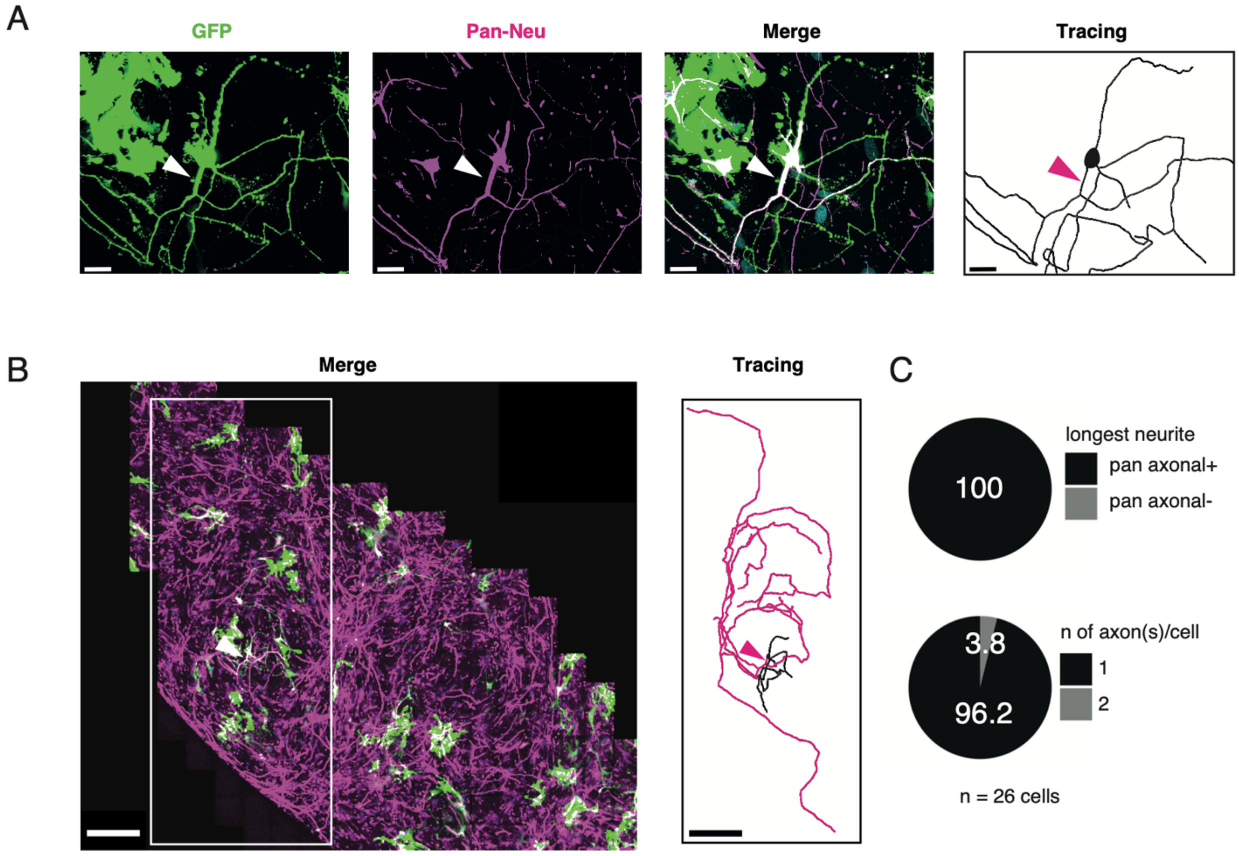
A
GFP
Pan-Neu
Merge
Tracing
B
Merge
Tracing
C
100
longest neurite
pan axonal+
pan axonal-
3.8
96.2
n of axon(s)/cell
1
2
n = 26 cells

**Supplementary Figure 4. Characterization of axons in iNs.** iNs were lipofected with a plasmid expressing cytosolic GFP (GFP, green), fixed at d35 and stained with an axonal marker (Pan-Neu, magenta). **A.** Chimpanzee iN stained with a pan-neurofilament antibody. The corresponding Imaris tracing is on the right. The arrowheads indicate that the longest of the neurites -as judged by the complete tracing in Imaris- is also positive for axonal marker (white arrowhead). Scale bar is 30 $\mu$m. **B**. Low magnification maximum intensity projection of GFP-labeled cells (green) stained with pan-neurofilament antibody (magenta). The Imaris tracing on the right corresponds to the white box. Magenta tracing corresponds to the longest neurite. Of note ape and human iNs generate long axons, compared to dendrites. As the longest neurite (axon) encompasses a large part of the total neurite length. Scale bar is 300 $\mu$m. **C**. Quantifications show that (i) for 100% of the stained cells, the longest neurite is positive for axonal marker (upper pie chart) (iii) the majority of the cell (96% of total) features one axon. 4% of the cells were found to have 2 axons (bottom pie chart). Of note, in our system the % of cells with two axons is lower than what reported for iNs autaptic culture [25]. This is possibly due to the NGN2-iNs system in conjunction with the culture conditions.

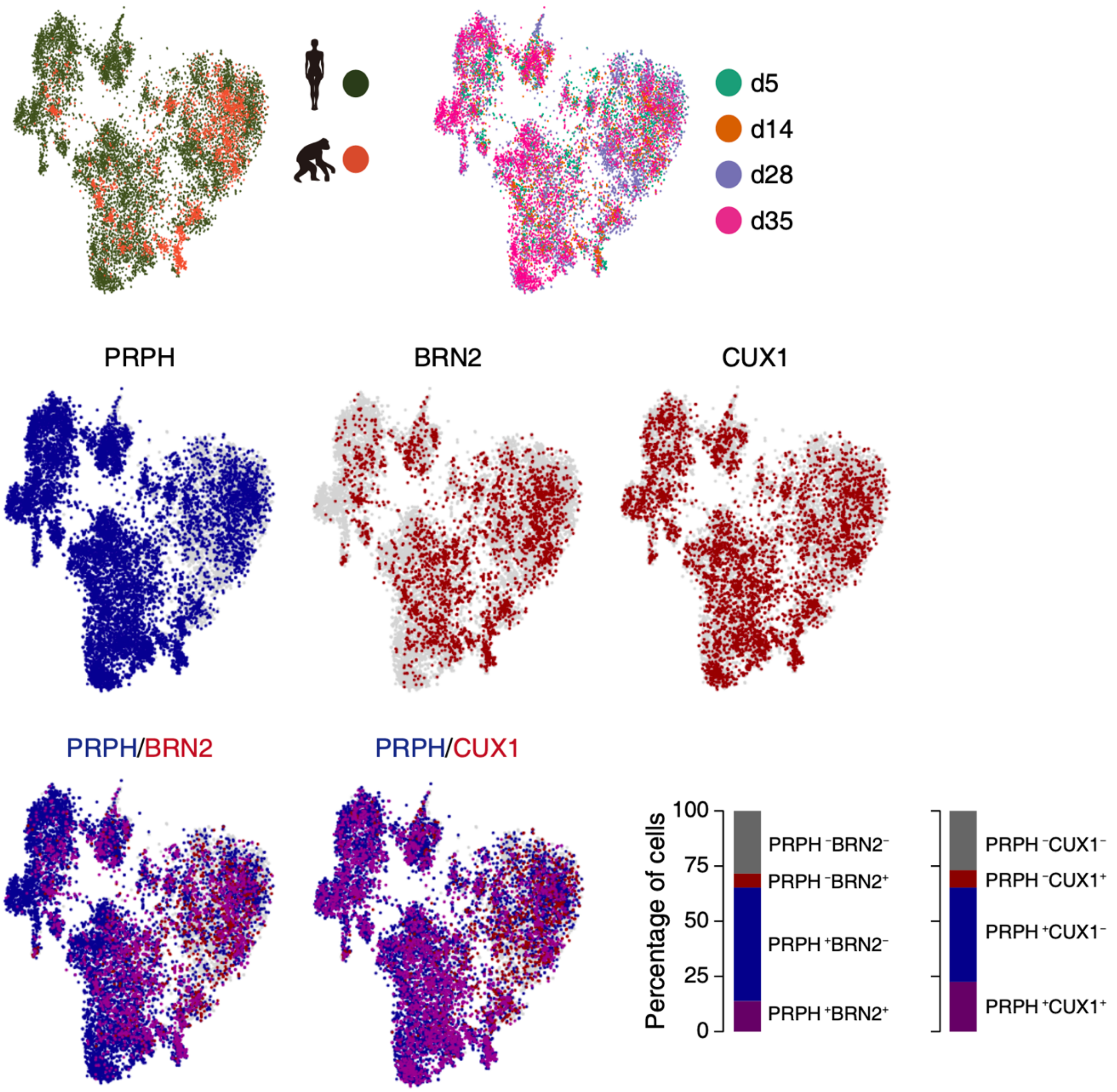
d5
d14
d28
d35
PRPH
BRN2
CUX1
PRPH/BRN2
PRPH/CUX1
Percentage of cells
100
75
50
25
0
PRPH⁻BRN2⁻
PRPH⁻BRN2⁺
PRPH⁺BRN2⁻
PRPH⁺BRN2⁺
PRPH⁻CUX1⁻
PRPH⁻CUX1⁺
PRPH⁺CUX1⁻
PRPH⁺CUX1⁺

**Supplementary Figure 5. iNs are sensory neurons.** tSNE plots showing single iNs from each species and at each timepoints (top panel). scRNAseq analysis reveals that NGN2-induced iNs express both sensory neuron markers as well as cortical markers. Cell identity and heterogeneity was assessed using tSNE combined for all time points (d5, d14, d28 and d35) and chimpanzee (SandraA) and human (409B2) cells together. Marker gene expression for single markers or a combination of sensory and cortical markers. Sensory neurons are shown in blue (*PRPH*) and cortical markers are shown in red (*BRN2* and *CUX1*). Quantifications show that around 65% of the neurons are *PRPH* positive, (i) 20% of neurons are *BRN2* positive and 13.8% of cells are *PRPH* and *BRN2* positive (left bar plot), (ii) 30% of cells are *CUX1* positive and 22.5% of cells are *PRPH* and *CUX1* positive (right bar plot), suggesting that the system consists of sensory and cortical neurons and possibly of cortical sensory neurons.

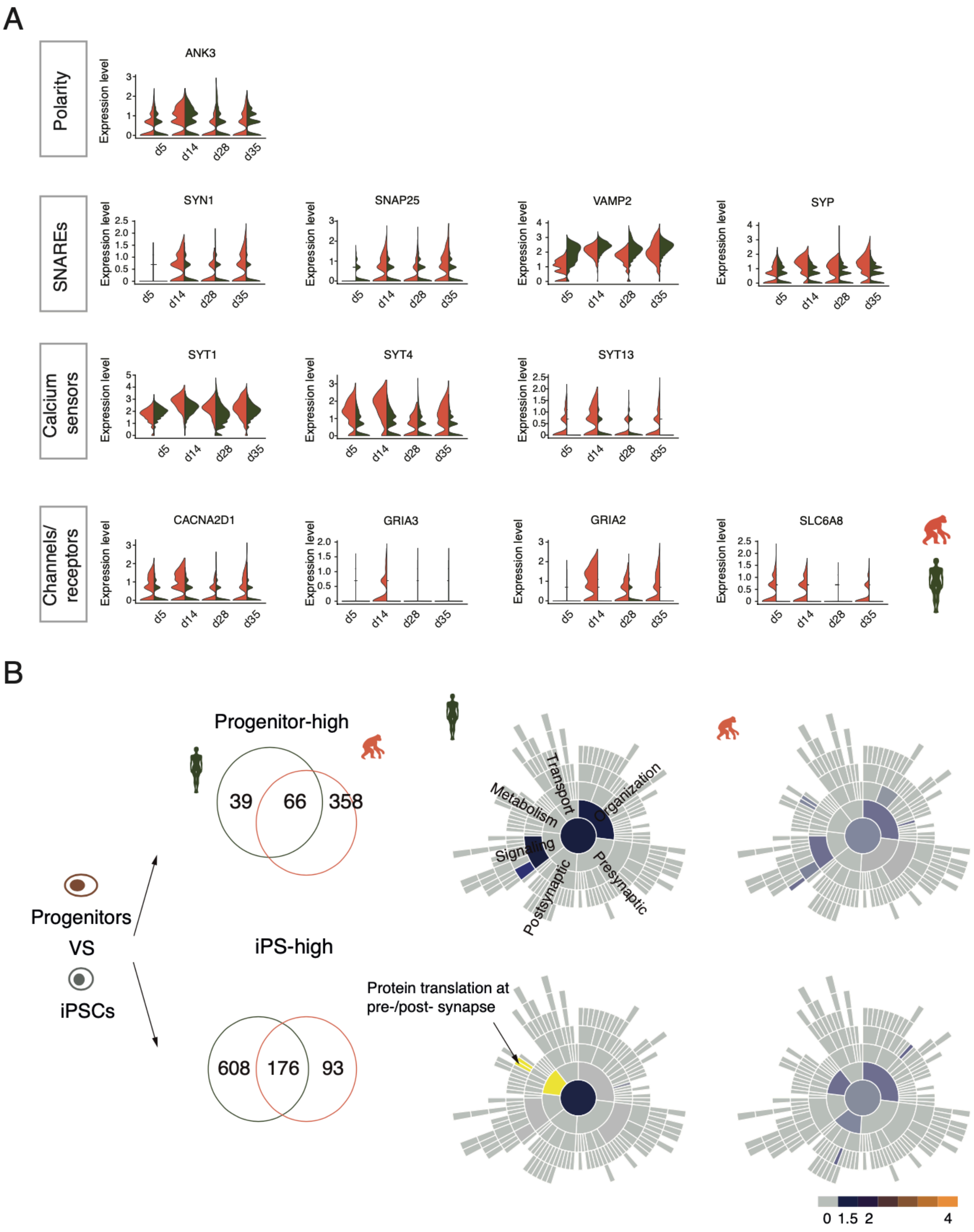

A
Polarity
ANK3
SNAREs
SYN1
SNAP25
VAMP2
SYP
Calcium sensors
SYT1
SYT4
SYT13
Channels/ receptors
CACNA2D1
GRIA3
GRIA2
SLC6A8
Expression level
d5
d14
d28
d35
B
Progenitors
VS
iPSCs
Progenitor-high
39
66
358
iPS-high
608
176
93
Transport
Organization
Metabolism
Signaling
Presynaptic
Postsynaptic
Protein translation at pre-/post- synapse
0
1.5
2
4

**Supplementary Figure 6. Transcriptional neoteny of synaptic genes in human iNs.**

**A.** Synaptic genes are differentially expressed in chimpanzee (SandraA) and human (409B2) iNs. Expression over time of a polarity gene (*ANK3*), SNARE proteins (*SNAP25*, *VAMP2* and the syntaxin-binding protein *SYP*), synapsin (*SYN1*), calcium sensors for regulated secretion (Synaptotagmin, *SYT1*, *4* and *13*) and channels/receptors (the voltage operated calcium channel's subunit *CACNA2D1*, the glutamate receptor's subunits *GRIA2* and *3* and the creatine transporter *SLC6A8*). **B.** SynGO enrichment analysis with significantly differentially expressed genes between NP and iPS cells of the same species. Left panel, higher expressed genes in human and chimpanzee NP (top panel) compared with corresponding iPS (bottom panel). Right panel, sunburst plots of enriched synaptic GO terms in human and chimpanzee NP and iPS cells. The log10-transformed FDR-corrected p value per ontology term is visualized for BP. Note that the bottom BP sunburst plots are aligned with the top-right one.

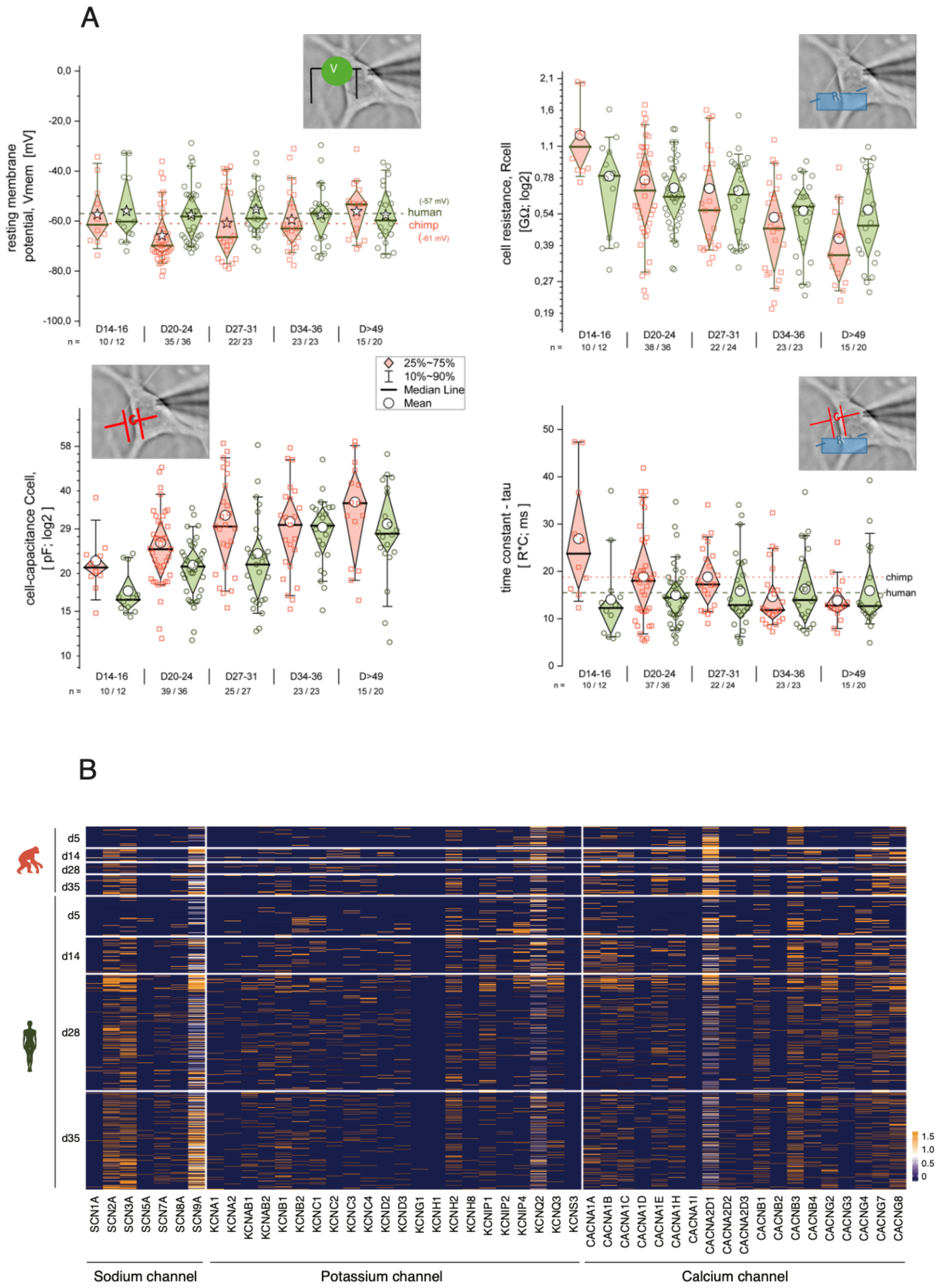
A
resting membrane potential, Vmem [mV]
0,0
-20,0
-40,0
-60,0
-80,0
-100,0
(-57 mV)
human
chimp
(-61 mV)
D14-16
D20-24
D27-31
D34-36
D>49
n =
10 / 12
35 / 36
22/ 23
23 / 23
15 / 20
cell resistance, Rcell [GΩ; log2]
2,1
1,6
1,1
0,78
0,54
0,39
0,27
0,19
10 / 12
38 / 36
22 / 24
23 / 23
15 / 20
25%~75%
10%~90%
Median Line
Mean
cell-capacitance Ccell, [ pF; log2 ]
58
40
29
20
15
10
10 / 12
39 / 36
25 / 27
23 / 23
15 / 20
time constant - tau [ R*C; ms ]
50
40
30
20
10
0
chimp
human
10 / 12
37 / 36
22 / 24
23 / 23
15 / 20
B
d5
d14
d28
d35
1.5
1.0
0.5
0
SCN1A SCN2A SCN3A SCN5A SCN7A SCN8A SCN9A
KCNA1 KCNA2 KCNAB1 KCNAB2 KCNB1 KCNB2 KCNC1 KCNC2 KCNC3 KCNC4 KCND2 KCND3 KCNG1 KCNH1 KCNH2 KCNH8 KCNIP1 KCNIP2 KCNIP4 KCNQ2 KCNQ3 KCNS3
CACNA1A CACNA1B CACNA1C CACNA1D CACNA1E CACNA1H CACNA1I CACNA2D1 CACNA2D2 CACNA2D3 CACNB1 CACNB2 CACNB3 CACNB4 CACNG2 CACNG3 CACNG4 CACNG7 CACNG8
Sodium channel
Potassium channel
Calcium channel

**Supplementary Figure 7. Basic electrical properties of apes and human iNs.**

Comparison of basic electrical properties between ape and human iNs 2 - 8 weeks post DOX-induced expression of NGN2 to verify the maturation of iNs. **A.** Basic electrical properties: Resting-membrane potential (Vrmp)**,** Cell-input resistances (Rcell), Cell capacitance (Ccell) and Time constant (tau = R*C; C; a measure of the cell response to stimulus). **B.** Heatmap showing the expression of voltage-gated sodium, potassium and calcium channels in chimpanzee (SandraA) and human (409B2) iNs.

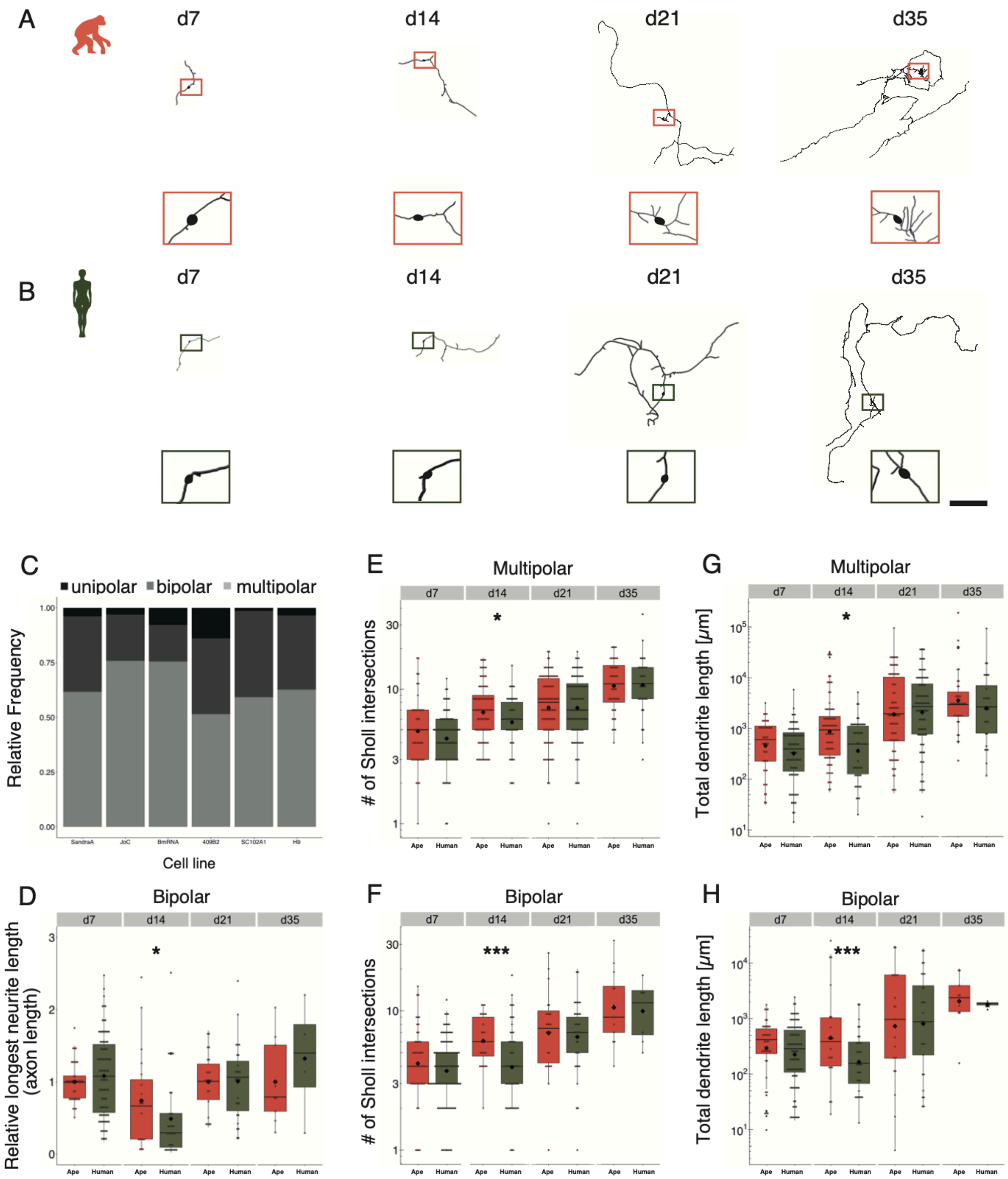
A
d7
d14
d21
d35
B
d7
d14
d21
d35
C
unipolar
bipolar
multipolar
Relative Frequency
Cell line
E
Multipolar
### of Sholl intersections
G
Multipolar
Total dendrite length [μm]
D
Bipolar
Relative longest neurite length (axon length)
F
Bipolar
### of Sholl intersections
H
Bipolar
Total dendrite length [μm]
Ape
Human

**Supplementary Figure 8. Dynamics of structural maturation in bipolar and multipolar iNs.** iNs were lipofected four days prior fixation with a plasmid expressing cytosolic GFP and fixed at different time points to quantify their structural features**.** **A, B**. Chimpanzee (**A**, SandraA, orange) and human (**B**, 409B2, green) bipolar iNs development over time. Zoom-ins show the cell body. Scale bar is 1 mm. **C**. Relative frequency of monopolar (black), bipolar (dark grey) and multipolar (grey) iNs for all the cell lines used in this study: chimpanzee (SandraA and JoC) and bonobo (BmRNA) and human (409B2, SC102A and H9). **D.** Relative longest neurite (axon) length in bipolar iNs. Ape iNs show a higher axon length at d14 ($p=0.0426$). Data representation as normalized data against the mean per batch of the ape data, the scale is logarithmic. **E, F**. Total number of Sholl intersections for multipolar (**E**) and bipolar (**F**) iNs. Apes iNs show a higher number of Sholl intersections at d14 for both multipolar iNs ($p=0.0469$) and bipolar ($p=0.0008$) iNs. The scale is logarithmic. **G, H.** Total dendrite length, expressed in microns for multipolar (**G**) and bipolar (**H**) iNs. Ape iNs show a higher total dendrite length at d14 for both multipolar ($p=0.0009$) and bipolar iNs ($p=0.04471$). The scale is logarithmic. For all graphs, black rhombs represent the mean, black lines the median. Significance score: $p<0.05$ *, $p<0.01$ **, $p<0.001$ ***, Mann-Whitney-U-Test for relative longest neurite length and number of Sholl intersections, Unpaired T-test for total dendrite length.
