## Supplementary Table 1 for "Comparison of induced neurons reveals slower structural and functional maturation in humans than in apes"

| | d7 | | | | d14 | | | | d21 | | | | d35 | | | |
|---|---|---|---|---|---|---|---|---|---|---|---|---|---|---|---|---|
| cellline/species | unipolar | bipolar | multipolar | total | unipolar | bipolar | multipolar | total | unipolar | bipolar | multipolar | total | unipolar | bipolar | multipolar | total |
| hiPS409-B2 | 18 | 34 | 24 | 76 | 2 | 10 | 21 | 33 | 1 | 9 | 29 | 39 | 1 | 4 | 11 | 16 |
| SC102A1 | 1 | 37 | 29 | 67 | 0 | 8 | 15 | 23 | 1 | 8 | 20 | 29 | 0 | 0 | 14 | 14 |
| H9 | 2 | 21 | 31 | 54 | 2 | 15 | 13 | 30 | 1 | 12 | 44 | 57 | 0 | 0 | 3 | 3 |
| **Human** | **21** | **92** | **84** | **197** | **4** | **33** | **49** | **86** | **3** | **29** | **93** | **125** | **1** | **4** | **28** | **33** |

| | d7 | | | | d14 | | | | d21 | | | | d35 | | | |
|---|---|---|---|---|---|---|---|---|---|---|---|---|---|---|---|---|
| cellline/species | unipolar | bipolar | multipolar | total | unipolar | bipolar | multipolar | total | unipolar | bipolar | multipolar | total | unipolar | bipolar | multipolar | total |
| SandraA | 4 | 30 | 26 | 60 | 0 | 9 | 19 | 28 | 1 | 9 | 27 | 38 | 1 | 5 | 16 | 22 |
| JoC | 2 | 6 | 6 | 14 | 0 | 7 | 31 | 38 | 1 | 6 | 24 | 31 | 0 | 1 | 10 | 11 |
| BmRNA | 3 | 2 | 13 | 18 | 1 | 5 | 28 | 34 | 2 | 7 | 30 | 39 | 0 | 3 | 6 | 9 |
| **Chimpanze Bonobo** | **9** | **38** | **45** | **92** | **1** | **21** | **78** | **100** | **4** | **22** | **81** | **108** | **1** | **9** | **32** | **42** |

Table 1. Schoernig et al.
