## Supplementary Table 2 for "Comparison of induced neurons reveals slower structural and functional maturation in humans than in apes"

Schoernig at al.

| REAGENT or RESOURCE | SOURCE | IDENTIFIER |
|---|---|---|
| **CONSUMABLES** | | |
| Cell culture test plates, sterile (6 well, 24 well) | TPP® | 92006, 92024 |
| Coverslips (pre-treated, 12 mm) | Kleinfeld Labortechnik | GG-12-Pre |
| Cryogenic vials | Thermo Fisher Scientific | 5000-0020 |
| Falcon tubes (15 ml, 50 ml) | Greiner bio-one | 188271, 210261 |
| Glass Pasteur pipettes | VWR | 612-1701 |
| Lens cleaning tissue | GE Healthcare Life Sciences | 2105-841 |
| Microscope slides | Thermo Fisher Scientific | J3800AMNZ |
| Parafilm | Carl Roth | PM-996 |
| Pipette tips, filter tips | Eppendorf, Gilson | |
| Precision wipes | Kimtech science | 7552 |
| Serological pipets, sterile (5 ml, 10 ml, 25 ml, 50 ml) | Corning® | |
| Single-use syringes (5 ml, 10 ml, 30 ml) | B Braun | |
| Sterile square media bottle (125 ml, 250 ml) | Nalgene | 2019-0125, 2019-0250 |
| Syringe-Filter (0.22 µm) | TPP® | 99722 |
| Tissue culture dish (40 mm) | TPP® | 93040 |
| Tissue culture flask, sterile (75 $cm^2$) | TPP® | 90076 |
| Tubes (0.5 ml, 1.5 ml, 2.0 ml) | Eppendorf | |
| **DEVICES** | | |
| Accu-jet® pro | Brand | |
| Analytical balance | Kern | AEJ-CM |
| Beaker (100 ml, 500 ml) | Schott Duran | |
| Bioanalyzer Instrument 2100 | Agilent | G2939Ba |
| Centrifuge | Thermo Fisher Scientific | Heraeus Megafuge 40R |
| Clean bench | Thermo Fisher Scientific | Heraeus Instruments |
| Confocal laser scanning microscope | OLYMPUS FV1200 | BX61W1 Multiphoton FV1000 Microscope |

| | | |
|---|---|---|
| | Fluorescent Light Source: | U-HGLGPS |
| | Power Supply: | FV10-MCPSU |
| Confocal laser scanning microscope | Zeiss LSM 780 NLO | Zeiss axio Examiner.Z1, upright stand |
| Countess™ automated cell counter | Invitrogen | |
| EPC-10 amplifier | HEKA, Lambrecht, Germany | |
| Freezing container | Nalgene | |
| Glass bottles (100 ml, 500 ml) | Schott Duran | |
| Glass electrodes for electrophysiology | Hilgenberg, Germany | 4.5-5.5 MΩ, oD/iD, borosilicate |
| HighSeq2500 | Ilumina | |
| Incubator Heracell 240 | Thermo Fisher Scientific | |
| Inverse microscope | Zeiss | Axiovert 200 |
| | MultiSpec-Micomager DualV | |
| | Filter Set: | 46HEYFP |
| Inverted microscope | Zeiss | Axio Observer.Z1 SMC200 |
| | | HXP12OV |
| | Fluorescent Light Source: | 232 |
| Magnetic stirring hot plate | Heidolph | MR3002 |
| Microcentrifuge | Carl Roth | |
| Nucleofector™2b | Lonza | |
| Pipettes (10 µl, 100 µl, 200 µl, 1000 µl) | Eppendorf, Gilson | |
| Spinning disc confocal microscope | Andor Revolution WD Borealis Mosaic | Andor IX 83, inverted stand |
| Stereo microscope | Olympus | SZX7 |
| Tweezers, Scoops | | |
| Upright microscope for electrophysiology | Olympus | BXW-51 |
| Vacusafe™ Vacuum aspiration system | Integra Biosciences | |
| Vortex-Genie 2 | Scientific Industries | |
| **REAGENTS** | | |
| Accutase® solution | Sigma Aldrich | A6964 |
| B27 Supplement (50x) | Gibco® | 17504-044 |
| BDNF, human | Promokine | C-66212 |
| Borax | Sigma Aldrich | B3545 |

| | | |
|---|---|---|
| Boric acid | Merck | 1.00165.1000 |
| $CaCl_2$ | Merck | |
| Chromium Single Cell 3' Library & Gel Bead Kit v2, 16 rxns | 10X Genomics | PN-120237 |
| Cytosine β-D-arabinofuranoside | Sigma Aldrich | C1768 |
| DAPI | Sigma Aldrich | D9542 |
| DMEM, high glucose | Gibco® | 11965-092 |
| DMEM/F-12 | Gibco® | 31330-038 |
| Dnase I | New England Biolabs | 2 U/µl |
| Doxycycline | Sigma Aldrich | D9891 |
| Dulbecco's Phosphate-Buffered Saline (DPBS) | Gibco® | 14190-094 |
| Ethylenediaminetetraacetic acid (EDTA) | Gibco® | 15575-038 |
| Ethanol | Carl Roth | K928.1 |
| Fetal bovine serum | Sigma Aldrich | F2442 |
| G418 disulfate salt solution | Sigma Aldrich | 68168 |
| Gelatine | Carl Roth | 4308.1 |
| Glucose | Sigma-Aldrich | 158968-500G |
| GlutaMAX | Gibco® | 35050-038 |
| Glycine | Serva | 23390.03 |
| HEPES | Sigma-Aldrich | H3375-250G |
| Hygromycin B-solution | Carl Roth | CP12.2 |
| Isopropanol | Merck | 1.09634.2511 |
| KCl | Merck | |
| K-gluconate | Merck | |
| Knockout DMEM/F12 | Gibco® | 12660-012 |
| Laminin | Sigma Aldrich | L2020 |
| Lipofectamine 3000 Transfection Kit | Invitrogen | C3000-008 |
| Matrigel Matrix | Corning® | 354277 |
| MEM non-essential amino acid solution (100x) | Sigma Aldrich | M7145 |
| mFreSR™ | Stemcell™ Technologies | 05853 |
| $MgCl_2$ | Merck | |
| Mowiol 4-88 | Sigma Aldrich | 81381 |
| mTeSR™1 | Stemcell™ Technologies | 85851 |
| mTeSR™1 5x Supplement | Stemcell™ Technologies | 85852 |

| | | |
|---|---|---|
| N2-Supplement (100x) | Thermo Fisher Scientific | 17502048 |
| NaCl | Merck | 1.06404.1000 |
| $NaHCO_3$ | Sigma-Aldrich | S5761-1KG |
| $Na_2HPO_4 \cdot 2\ H_2O$ | Merck | 1.06580.1000 |
| $NaH_2PO_4 \cdot H_2O$ | Merck | 1.06346.1000 |
| NaOH | Merck | 1.06462.1000 |
| Neurobasal™ Medium | Gibco® | 21103-049 |
| NT3, human | Promokine | C-66425 |
| Opti-MEM™ | Gibco® | 31985-070 |
| Paraformaldehyde, 96 % | ACROS Organics™ | 30525-89-4 |
| Pen/Strep | Gibco® | 15140-122 |
| Percoll | Sigma-Aldrich | P1644-100ML |
| Phosphate-Buffered Saline (PBS) | Gibco® | 10010-015 |
| Poly-D-Lysine solution | Sigma Aldrich | A-003-E |
| Puromycin dihydrochloride | Sigma Aldrich | P9620 |
| Rock-Inhibitor Y-27632 | Stemcell™ Technologies | 72305 |
| Sucrose | Merck | 1.07653.100 |
| Triton X-100 Solution | Sigma Aldrich | 93443 |
| TrypLE™ Express (1x) | Gibco® | 12605-010 |
| UltraPure™ Destilled Water | Invitrogen | 10977-035 |
| **ANTIBODIES** | | |
| Anti MAP2 Antibody | Invitrogen | PA1-16751<br>Dilution 1:1000 |
| Anti-Synapsin ½ guineapig antiserum | Synaptic Systems | 106004<br>Dilution 1:1000 |
| Purified anti-Tubulinβ3 (TUBB3), Clone: TUJ1<br>Coupled to AlexaFlour™ 488 or Alexa Flour™ 555 | BioLegend | 801202<br>Dilution 1:1000 |
| Purified anit-Neurofilament Marker (pan-axonal, cocktail), Clone: SMI312 | BioLegend | 8379074<br>Dilution 1:400 |
| Donkey anti-Mouse IgG (H+L) Highly Cross-Adsorbed Secondary Antibody, Alexa Fluor™ 555 | Invitrogen | A21202<br>Dilution 1:1000 |

| | | |
|---|---|---|
| Goat anti-Rabbit IgG (H+L) Highly Cross-Adsorbed Secondary Antibody, Alexa Fluor$^{TM}$ Plus 555 | Invitrogen | A32732<br>Dilution 1:1000 |
| Goat anti-Chicken IgG Highly Cross-Adsorbed Secondary Antibody, Alexa Fluor$^{TM}$ 488 | Invitrogen | A11039<br>Dilution 1:1000 |
| Goat anti-Guinea Pig IgG Highly Cross-Adsorbed Secondary Antibody, Alexa FluorTM 568 | Invitrogen | A11075<br>Dilution 1:1000 |
| **CELLS** | | |
| Human | 409B2 iPS cell line<br>409B2_Ngn2 | Riken BRC Cellbank<br>Generated by Maria Schörnig |
| Human | SC102A-1 iPS cell line<br>SC102A-1_Ngn2 | Systems Biosciences<br>Generated by Maria Schörnig |
| Human | HmRNA iPS cell line<br>HmRNA_Ngn2 | Generated by Anne Weigert<br>Generated by Anne Weigert |
| Human | H9 ES cell line<br>H9_Ngn2 | WiCell<br>Generated by Maria Schörnig |
| Chimpanzee | Sandra A<br>Sandra A_Ngn2 | Generated in a previous study.<br>Generated by Maria Schörnig |
| Chimpanzee | Jo_C<br>Jo_C_Ngn2 | Generated in a previous study.<br>Generated by Maria Schörnig |
| Chimpanzee | ciPS01<br>Chimp male iPSC Sendai CL5<br><br><br>ciPS01_Ngn2 | Provided by the Max-Delbrück-Centrum für Molekulare Medizin<br>Generated by Anne Weigert. |
| Bonobo | BmRNA<br>BmRNA_Ngn2 | Reprogramming by mRNA<br>Generated by Maria Schörnig |
| Primary Rat Astrocytes | RjHan:WI – Wistar rat from Janvier | Primary cortical rat astrocytes were freshly prepared for this study |
| Human dermal fibroblasts | Lonza | CC-2511 |
| **PLASMIDS** | | |
| pCMV(CAT)T7-SB100 | Addgene | 34879 |

| | | |
|---|---|---|
| pmax GFP | Lonza | D-00072 |
| pSBbi-RH | Addgene | 60516 |
| **SOFTWARE** | | |
| AxioVision Rel. | 4.8 | |
| FV10-ASW | 4.2 | |
| Image J | v1.51w. | |
| Imaris | 9.5. | |
| Imaris File Converter | 9.2.0 | |
| Imaris Stitcher | 9.2.0 | |
| Origin | OriginLab version 2018-2019b | |
| Patch-and Fitmaster software | HEKA version 2.9x | |
| R | 3.5.1 | |
| ZEN | | |
| **SOLUTIONS, BUFFERS AND MEDIA** | | |
| PFA 4 % | 4% PFA<br>4% Sucrose<br>120 mM sodium phosphate buffer pH 7.4 | |
| Sodium phosphate buffer (240mM) | 240 mM $Na_2HPO_4$<br>240 mM $NaH_2PO_4$, pH 7.4 | |
| Astrocyte medium | DMEM high glucose<br>10% FBS<br>1% Pen/Strep | |
| Borate-Buffer, pH 8.4 | boric acid<br>borax | |
| Immunofluorescence buffer (IF buffer) | 30 mM NaCL<br>0.2% gelatine<br>0.05% Triton X-100<br>120 mM phosphate buffer | |
| Glycine buffer | 0.2 M glycine<br>120 mM phosphate buffer | |

| | | |
|---|---|---|
| Permeabilization buffer | 0.05% Triton X-100<br>120 mM phosphate buffer | |
| Artificial cerebrospinal fluid | 100 mM NaCl<br>3.5 mM KCl<br>1 mM $MgCl_2$<br>2 mM $CaCl_2$<br>30 mM $NaHCO_3$<br>1.25 mM $NaH_2PO_4$<br>10 mM glucose | |
| Internal solution electrophysiology | 130 mM K-gluconate<br>10 mM NaCl<br>4 mM Mg-ATP<br>0.5 GTP<br>10 mM HEPES<br>0.05 mM EGTA | |
| **KITs** | | |
| Stem MACS mRNA transfection Kit | Miltenyi Biotec | 130-104-463 |
| Human Pluripotent Stem Cell 3 Colour Immunohistochemistry Kit | R&D Systems1 | SC021 |
| Human Pluripotent Stem Cell Functional Identification Kit | R&D Systems | SC027B |
| StemMACS Trilineage Differentiation Kit | Miltenyi Biotec | 130-115-660 |
