## Supplementary Table 3 for "Comparison of induced neurons reveals slower structural and functional maturation in humans than in apes"

**Table 1: Statistics on basic electrical properties (Vmem, Rcell, Ccell).** Shown is for different days of differentiation and species the n number of cells analyzed, the median, mean, SD and SMEM values for Vrmp, Rcell and Ccell and P values for a 2-Way an anova analyses on age and species effects and interaction of species and age. Bold values represent significant p values.

| **day** | **species** | **Vrmp** | **Vrmp** | **Vrmp** | **N** | **p-val.** | **Rcell** | **Rcell** | **Rcell** | **N** | **p** | **Ccell** | **Ccell** | **Ccell** | **N** | **p** |
|---|---|---|---|---|---|---|---|---|---|---|---|---|---|---|---|---|
| days | | mV median | mV mean | mV SEM | # | days spec*days | MOhm median | MOhm mean | MOhm SEM | # | days spec*days | pF median | pF mean | pF SEM | # | days spec*days |
| D14-16 | chimp/bonobo | -61 | -57 | 4,1 | 10 | $F_{days}$ 1,4; p = 0,23 | 1068 | 1200 | 154 | 10 | **$F_{days}$ 11,7; p = 1,3 E-8** | 21 | 22 | 2,0 | 10 | **$F_{days}$ 10; p = 1,8 E-7** |
| | human | -60 | -56 | 3,8 | 12 | | 793 | 789 | 110 | 12 | | 16 | 17 | 1,0 | 12 | |
| D20-24 | chimp/bonobo | -70 | -66 | 1,9 | 35 | | 682 | 760 | 60 | 38 | | 25 | 26 | 1,4 | 39 | |
| | human | -58 | -58 | 1,9 | 36 | | 640 | 699 | 43 | 36 | | 21 | 21 | 0,9 | 36 | |
| D27-31 | chimp/bonobo | -66 | -61 | 3,1 | 22 | | 557 | 696 | 81 | 22 | | 30 | 33 | 2,7 | 25 | |
| | human | -59 | -55 | 2,2 | 23 | | 655 | 680 | 61 | 24 | | 22 | 24 | 2,0 | 27 | |
| D34-36 | chimp/bonobo | -63 | -59 | 2,7 | 23 | | 462 | 518 | 53 | 23 | | 30 | 31 | 2,5 | 23 | |
| | human | -57 | -58 | 2,5 | 23 | | 578 | 554 | 43 | 23 | | 30 | 29 | 1,8 | 23 | |
| D>49 | chimp/bonobo | -53 | -56 | 2,9 | 15 | | 352 | 416 | 46 | 15 | | 36 | 36 | 3,6 | 15 | |
| | human | -60 | -58 | 2,7 | 20 | | 477 | 559 | 60 | 20 | | 28 | 30 | 2,5 | 20 | |
| p val. (species / spec.*days) | | $F_{species}$ 3,1; p = 0,081 | | | | $F_{spec*days}$ 1,2; p = 0,33 | $F_{species}$ 1,9; p = 0,17 | | | | **$F_{spec*days}$ 2,9; p = 0,024** | **$F_{species}$ 13; p = 3,2 E-4** | | | | $F_{spec*days}$ 0,9; p = 0,45 |

**Table 2: Statistics on basic electrical properties (tau, APs and sEPSCs).** Shown is for different days of differentiation and species the n number of cells analyzed, the median, mean, SD and SMEM values for tau, APs and sEPSCs and P values for an unpaired t-test with welch correction for pairwise comparison of APs and EPSCs between the species and P values for an anova analyses on age and species effects and interaction of species and age. Bold values represent significant p values.

| day | species | Tau | Tau | Tau | N | p | APs | APs | APs | N | p | p | EPSCs | EPSCs | EPSCs | N | p | p |
|---|---|---|---|---|---|---|---|---|---|---|---|---|---|---|---|---|---|---|
| days | | ms median | ms mean | ms SEM | # | days spec*days | # median | # mean | # SEM | # | | days spec*days | Hz median | Hz mean | Hz SEM | # | | days spec*days |
| D14-16 | ape | 23,7 | 26,9 | 12,9 | 10 | $F_{days}$ 1,9; p = 0,11 | 10 | 10 | 2 | 10 | **0,021** | **$F_{days}$ 4,46; p = 0,00175** | 0,30 | 0,27 | 0,025 | 11 | **0,045** | **$F_{days}$ 12; p = 2,8E-09** |
| | human | 12,3 | 14,0 | 9,27 | 12 | | 1 | 4 | 1 | 13 | | | 0,05 | 0,08 | 0,024 | 16 | | |
| D20-24 | ape | 18,0 | 18,9 | 10,1 | 37 | | 12 | 11 | 1 | 39 | **0,0002** | | 2,91 | 1,39 | 0,024 | 52 | **1,5E-07** | |
| | human | 14,4 | 15,0 | 6,70 | 36 | | 5 | 6 | 1 | 36 | | | 0,12 | 0,14 | 0,021 | 51 | | |
| D27-31 | ape | 17,2 | 18,8 | 6,72 | 22 | | 12 | 12 | 1 | 25 | **0,0011** | | 4,73 | 6,96 | 0,021 | 25 | **3,8E-07** | |
| | human | 12,9 | 15,8 | 8,29 | 24 | | 7 | 7 | 1 | 27 | | | 0,28 | 0,34 | 0,026 | 30 | | |
| D34-36 | ape | 11,8 | 14,6 | 6,58 | 23 | | 13 | 12 | 1 | 23 | **0,0085** | | 12,42 | 5,55 | 0,027 | 31 | **0,019** | |
| | human | 14,0 | 16,3 | 8,20 | 23 | | 7 | 8 | 1 | 23 | | | 0,52 | 1,10 | 0,026 | 30 | | |
| D>49 | ape | 12,8 | 13,8 | 4,86 | 15 | | 15 | 13 | 2 | 15 | 0,54 | | 9,38 | 3,03 | 0,028 | 20 | 0,22 | |
| | human | 12,7 | 15,9 | 8,66 | 20 | | 14 | 12 | 2 | 20 | | | 2,13 | 1,25 | 0,027 | 26 | | |
| p val. (species / spec.*days) | | **$F_{species}$ 7,2; p = 0,0080** | | | | **$F_{spec*days}$3,7; p = 0,0063** | **$F_{species}$ 29,1; p = 1,73E-07** | | | | | $F_{spec*days}$ 0,749; p = 0,559 | **$F_{species}$ 41; p = 7,3E-10** | | | | | $F_{spec*days}$ 2,0; p = 0,093 |
