## Supplementary Table 4 for "Comparison of induced neurons reveals slower structural and functional maturation in humans than in apes"

**Table 1: Statistics total neurite length of bipolar iNs.** Shown is for different days of differentiation and species the n number of cells analyzed, the differences between species in %, the mean total neurite length [$\mu$m], the sd [$\mu$m] and the p value for a pairwise comparison (Mann-Whitney-U-Test). The differences in % between the species are indicated with "-" for a difference where ape iNeurons show higher values and "+" for differences where the human iNs show higher values. Additionally, we show the values for a shapiro test on total neurite length and a 2-way anova on the interaction of species and day. Significant p values are shown in bold.

| Days of differentiation | Species | N | Differences in % | Mean [$\mu$m] | SD [$\mu$m] | P value (species) |
|---|---|---|---|---|---|---|
| **d7** | ape | 38 | -13% | 1916 | 791 | **0.0278** |
| | human | 92 | | 1673 | 1056 | * |
| **d14** | ape | 21 | -82% | 11184 | 17897 | **0.0003** |
| | human | 33 | | 1960 | 1580 | *** |
| **d21** | ape | 22 | -8% | 16699 | 16947 | 0.4508 |
| | human | 29 | | 15415 | 14553 | |
| **d35** | ape | 9 | +1% | 46158 | 49394 | 0.4693 |
| | human | 4 | | 46839 | 38651 | |
| **Spariro test** | **W** | **P value** | | | | |
| | 0.89869 | 7.215e-12 | | | | |
| **2 Way Anova** | **Df** | **Sum Sq** | **Mean Sq** | **F value** | **P value** | |
| **species** | 1 | 5.447 | 5.4466 | 35.6988 | **8.262e-09** | *** |
| **day** | 3 | 31.325 | 10.4416 | 68.4379 | **<2.2e-16** | *** |
| **Species*day** | 3 | 2.070 | 0.6900 | 4.522 | **0.004176** | ** |
| **Residual** | 240 | 1661.75 | 6.924 | | | |

**Table 2: Statistics total neurite length of multipolar iNs.** Shown is for different days of differentiation and species the n number of cells analyzed, the differences between species in %, the mean total neurite length [$\mu$m], the sd [$\mu$m] and the p value for a pairwise comparison (Mann-Whitney-U-Test). The differences in % between the species are indicated with "-" for a difference where ape iNeurons show higher values and "+" for differences where the human iNs show higher values. Additionally, we show the values for a shapiro test on total neurite length and a 2-way anova on the interaction of species and day. Significant p values are shown in bold.

| Days of differentiation | Species | N | Differences in % | Mean [$\mu$m] | SD [$\mu$m] | P value (species) |
|---|---|---|---|---|---|---|
| **d7** | ape | 45 | -19% | 2353 | 1160 | **0.0098** |
| | human | 84 | | 1905 | 1137 | ** |
| **d14** | ape | 78 | -49% | 8082 | 13560 | **0.0215** |
| | human | 49 | | 4093 | 5848 | * |
| **d21** | ape | 93 | +10% | 22594 | 23493 | 0.1434 |
| | human | 81 | | 24884 | 22216 | |
| **d35** | ape | 40 | -6% | 35615 | 45846 | 0.4088 |
| | human | 28 | | 33325 | 40643 | |
| **Spariro test** | **W** | **p-value** | | | | |
| | 0.95237 | 1.384e-11 | | | | |
| **2 Way Anova** | **Df** | **Sum Sq** | **Mean Sq** | **F value** | **P value** | |
| **species** | 1 | 0.696 | 0.6958 | 3.1951 | 0.0749 | |
| **day** | 3 | 77.186 | 25.7287 | 118.1547 | **<2e-16** | *** |
| **Species*day** | 3 | 1.561 | 0.5202 | 2.3889 | 0.0681 | |
| **Residual** | 490 | 106.700 | 0.278 | | | |

**Table 3: Statistics relative total neurite (axon) length of multipolar iNs.** Shown is for different days of differentiation and species the n number of cells analyzed, the differences between species in %, the mean relative longest neurite length [$\mu$m] corrected for batch effects, the sd [$\mu$m] and the p value for a pairwise comparison (Mann-Whitney-U-Test). The differences in % between the species are indicated with "-" for a difference where ape iNeurons show higher values and "+" for differences where the human iNs show higher values. Additionally, we show the values for a shapiro test on relative longest neurite (axon) length and a 2-way anova on the interaction of species and day. Significant p values are shown in bold.

| Days of differentiation | Species | N | Differences in % | Mean [$\mu$m] | SD [$\mu$m] | P value (species) |
|---|---|---|---|---|---|---|
| **d7** | ape | 45 | -13% | 1242 | 543 | 0.3327 |
| | human | 84 | | 1073 | 541 | |
| **d14** | ape | 78 | -28% | 3436 | 4621 | **0.0294** |
| | human | 49 | | 2461 | 4033 | * |
| **d21** | ape | 93 | +20% | 10008 | 9040 | 0.0587 |
| | human | 81 | | 12045 | 9948 | |
| **d35** | ape | 40 | +25% | 12271 | 11318 | **0.0094** |
| | human | 28 | | 15330 | 12689 | ** |
| **Spariro test** | **W** | **p-value** | | | | |
| | 0.71661 | 2.2e-16 | | | | |
| **2 Way Anova** | **Df** | **Sum Sq** | **Mean Sq** | **F value** | **P value** | |
| **species** | 1 | 3.46 | 3.4634 | 4.5319 | **0.0338** | * |
| **day** | 17 | 19.31 | 1.1356 | 1.4859 | 0.0949 | |
| **Species*day** | 17 | 20.19 | 1.1877 | 1.5541 | 0.0726 | |
| **Residual** | 462 | 353.08 | 0.7642 | | | |

**Table 4: Statistics relative longest neurite (axon) length of bipolar iNs.** Shown is for different days of differentiation and species the n number of cells analyzed, the differences between species in %, the mean relative longest neurite length [$\mu$m] corrected for batch effects, the sd [$\mu$m] and the p value for a pairwise comparison (Mann-Whitney-U-Test). The differences in % between the species are indicated with "-" for a difference where ape iNeurons show higher values and "+" for differences where the human iNs show higher values. Additionally, we show the values for a shapiro test on relative longest neurite (axon) length and a 2-way anova on the interaction of species and day. Significant p values are shown in bold.

| Days of differentiation | Species | N | Differences in % | Mean [$\mu$m] | SD [$\mu$m] | P value (species) |
|---|---|---|---|---|---|---|
| **d7** | ape | 38 | +11% | 1192 | 529 | 0.2763 |
| | human | 92 | | 1323 | 705 | |
| **d14** | ape | 21 | -51% | 6859 | 11701 | **0.0426** |
| | human | 33 | | 3347 | 969 | * |
| **d21** | ape | 22 | +38% | 8665 | 9518 | 0.3413 |
| | human | 29 | | 11958 | 7464 | |
| **d35** | ape | 9 | +32% | 18356 | 10778 | 0.2437 |
| | human | 4 | | 24274 | 14934 | |
| **Spariro test** | **W** | **P value** | | | | |
| | 0.7002 | 2.2e-16 | | | | |
| **2 Way Anova** | **Df** | **Sum Sq** | **Mean Sq** | **F value** | **P value** | |
| **species** | 1 | 0.061 | 0.06088 | 0.1177 | 0.7319 | |
| **day** | 13 | 30.318 | 2.33219 | 4.5076 | **8.280e-07** | *** |
| **Species*day** | 13 | 28.536 | 2.19505 | 4.2423 | **2.531e-06** | *** |
| **Residual** | 220 | 113.826 | 0.51739 | | | |

**Table 5: Statistics relative number of sholl intersections of multipolar iNs.** Shown is for different days of differentiation and species the n number of cells analyzed, the differences between species in %, the mean number of sholl intersections, the sd and the p value for a pairwise comparison (Mann-Whitney-U-Test). The differences in % between the species are indicated with "-" for a difference where ape iNeurons show higher values and "+" for differences where the human iNs show higher values. Additionally, we show the values for a shapiro test on number of sholl intersections and a 2-way anova on the interaction of species and day. Significant p values are shown in bold.

| **Days of differentiation** | **Species** | **N** | **Differences in %** | **Mean** | **SD** | **P value (species)** |
|---|---|---|---|---|---|---|
| **d7** | ape | 45 | -18% | 5.7500 | 3.5937 | 0.1419 |
| | human | 84 | | 4.7349 | 2.0487 | |
| **d14** | ape | 78 | -17% | 7.5402 | 3.6593 | **0.0469** * |
| | human | 49 | | 6.2444 | 2.6642 | |
| **d21** | ape | 93 | -1% | 8.3438 | 4.0570 | 0.4445 |
| | human | 81 | | 8.2577 | 3.9430 | |
| **d35** | ape | 32 | +4% | 11.5455 | 4.6976 | 0.5414 |
| | human | 28 | | 12.0000 | 6.0245 | |
| **Shapiro test** | **W** | **P value** | | | | |
| | 0.97969 | 8.769e-07 | | | | |
| **2 Way Anova** | **Df** | **Sum Sq** | **Mean Sq** | **F value** | **P value** | |
| **species** | 1 | 0.3936 | 0.39359 | 8.0897 | **0.0046** | ** |
| **day** | 3 | 7.1105 | 2.37016 | 48.7148 | **<2.2e-16** | *** |
| **Species*day** | 3 | 0.1441 | 0.04803 | 0.9872 | 0.3984 | |
| **Residual** | 527 | 25.6405 | 0.04865 | | | |

**Table 6: Statistics relative number of sholl intersections of bipolar iNs.** Shown is for different days of differentiation and species the n number of cells analyzed, the differences between species in %, the mean number of sholl intersections, the sd and the p value for a pairwise comparison (Mann-Whitney-U-Test). The differences in % between the species are indicated with "-" for a difference where ape iNeurons show higher values and "+" for differences where the human iNs show higher values. Additionally, we show the values for a shapiro test on number of sholl intersections and a 2-way anova on the interaction of species and day. Significant p values are shown in bold.

| **Days of differentiation** | **Species** | **N** | **Differences in %** | **Mean** | **SD** | **P value (species)** |
|---|---|---|---|---|---|---|
| **d7** | ape | 38 | -14% | 4.9608 | 2.9797 | 0.1043 |
| | human | 92 | | 4.2456 | 2.1351 | |
| **d14** | ape | 21 | -26% | 6.5714 | 2.5593 | **0.0008** *** |
| | human | 33 | | 4.8679 | 3.4027 | |
| **d21** | ape | 22 | -10% | 8.3846 | 5.4851 | 0.4256 |
| | human | 29 | | 7.6176 | 4.1048 | |
| **d35** | ape | 9 | -13% | 12.4615 | 7.9332 | 0.4137 |
| | human | 4 | | 10.875 | 4.6117 | |
| **Spariro test** | **W** | **P value** | | | | |
| | 0.97347 | 1.011e-05 | | | | |
| **2 Way Anova** | **Df** | **Sum Sq** | **Mean Sq** | **F value** | **P value** | |
| **species** | 1 | 1.1079 | 1.10787 | 17.0543 | **4.644e-05** | *** |
| **day** | 3 | 4.4735 | 1.49118 | 22.9548 | **1.761e-13** | *** |
| **Species*day** | 3 | 3.0.2924 | 0.09746 | 1.5003 | 0.2144 | |
| **Residual** | 319 | 20.7227 | 0.06496 | | | |

**Table 7: Statistics total dendrite length of multipolar iNs.** Shown is for different days of differentiation and species the n number of cells analyzed, the differences between species in %, the mean total dendrite length [$\mu$m], the sd [$\mu$m] and the p value for a pairwise comparison (Mann-Whitney-U-Test). The differences in % between the species are indicated with "-" for a difference where ape iNeurons show higher values and "+" for differences where the human iNs show higher values. Additionally, we show the values for a shapiro test on number of total dendrite length and a 2-way anova on the interaction of species and day. Significant p values are shown in bold.

| Days of differentiation | Species | N | Differences in % | Mean [$\mu$m] | SD [$\mu$m] | P value (species) |
|---|---|---|---|---|---|---|
| **d7** | ape | 45 | -20% | 776 | 101 | 0.1226 |
| | human | 84 | | 618 | 84.4 | |
| **d14** | ape | 78 | -72% | 2740 | 666 | **0.0009** *** |
| | human | 49 | | 770 | 136 | |
| **d21** | ape | 93 | -12% | 6843 | 1406 | 0.7268 |
| | human | 81 | | 6001 | 905 | |
| **d35** | ape | 32 | -41% | 13356 | 6113 | 0.3336 |
| | human | 28 | | 7926 | 3328 | |
| **Shapiro test** | **W** | **P value** | | | | |
| | 0.99504 | 0.1111 | | | | |
| **2 Way Anova** | **Df** | **Sum Sq** | **Mean Sq** | **F value** | **P value** | |
| **species** | 1 | 4.331 | 4.3312 | 10.3340 | **0.0014** | ** |
| **day** | 3 | 59.997 | 19.9989 | 47.7157 | **<2.2e-16** | *** |
| **Species*day** | 3 | 3.074 | 1.0248 | 2.4451 | 0.0633 | |
| **Residual** | 490 | 205.372 | 0.4191 | | | |

**Tabelle 8: Statistics total dendrite length of bipolar iNs.** Shown is for different days of differentiation and species the n number of cells analyzed, the differences between species in %, the mean total dendrite length [$\mu$m], the sd [$\mu$m] and the p value for a pairwise comparison (Mann-Whitney-U-Test). The differences in % between the species are indicated with "-" for a difference where ape iNeurons show higher values and "+" for differences where the human iNs show higher values. Additionally, we show the values for a shapiro test on number of total dendrite length and a 2-way anova on the interaction of species and day. Significant p values are shown in bold.

| Days of differentiation | Species | N | Differences in % | Mean | SD | P value (species) |
|---|---|---|---|---|---|---|
| **d7** | ape | 38 | -13% | 505 | 71.8 | 0.2874 |
| | human | 92 | | 440 | 49.2 | |
| **d14** | ape | 21 | -89% | 2918 | 1404 | **0.0447** * |
| | human | 33 | | 332 | 77.2 | |
| **d21** | ape | 22 | -11% | 3630 | 1214 | 0.8698 |
| | human | 29 | | 3234 | 896 | |
| **d35** | ape | 9 | -43% | 3175 | 888 | 0.7489 |
| | human | 4 | | 1821 | 129 | |
| **Spariro test** | **W** | **P value** | | | | |
| | 0.99165 | 0.1712 | | | | |
| **2 Way Anova** | **Df** | **Sum Sq** | **Mean Sq** | **F value** | **P value** | |
| **species** | 1 | 3.403 | 3.4033 | 7.4732 | **0.006729** | ** |
| **day** | 3 | 15.794 | 5.2648 | 11.5608 | **4.184e-07** | *** |
| **Species*day** | 3 | 3.1.552+08 | 0.5172 | 1.1358 | 0.335257 | |
| **Residual** | 240 | 109.296 | 0.04554 | | | |
